# Type I Interferon acts as a major barrier to the establishment of infectious bursal disease virus (IBDV) persistent infections

**DOI:** 10.1101/2020.10.09.333161

**Authors:** Laura Broto, Nicolás Romero, Fernando Méndez, Elisabet Diaz-Beneitez, Oscar Candelas-Rivera, Daniel Fuentes, Liliana L. Cubas-Gaona, Céline Courtillon, Nicolas Eterradosi, Sébastien M. Soubies, Juan R. Rodríguez, Dolores Rodríguez, José F. Rodríguez

**Affiliations:** Departamento de Biología Molecular y Celular, Centro Nacional de Biotecnología-CSIC, Madrid, Spain; OIE Reference Laboratory for Gumboro Disease, Avian and Rabbit Virology Immunology and Parasitology Unit (VIPAC), French Agency for Food, Environmental and Occupational Heath Safety (ANSES), Ploufragan, France

## Abstract

Infectious bursal disease virus (IBDV), the best characterized member of the *Birnaviridae* family, is a highly relevant avian pathogen causing both acute and persistent infections in different avian hosts. Here, we describe the establishment of clonal, long-term, productive persistent IBDV infections in DF-1 chicken embryonic fibroblasts. Although virus yields in persistently-infected cells are exceedingly lower than those detected in acutely infected cells, the replication fitness of viruses isolated from persistently-infected cells is higher than that of the parental virus. Persistently-infected DF-1 and IBDV-cured cell lines derived from them do not respond to type I interferon (IFN). High-throughput genome sequencing revealed that this defect is due to mutations affecting the IFNα/β receptor subunit 2 (IFNAR2) gene resulting in the expression of IFNAR2 polypeptides harbouring large C-terminal deletions that abolish the signalling capacity of IFNα/β receptor complex. Ectopic expression of a recombinant chicken IFNAR2 gene efficiently rescues IFNα responsiveness. IBDV-cured cell lines derived from persistently infected cells exhibit a drastically enhanced proneness to establishing new persistent IBDV infections. Additionally, experiments carried out with human HeLa cells lacking the IFNAR2 gene fully recapitulate results obtained with DF-1 cells, exhibiting a highly enhanced capacity to both survive the acute IBDV infection phase and to support the establishment of persistent IBDV infections. Results presented here show that the inactivation of the JAK-STAT signalling pathway significantly reduces the apoptotic response induced by the infection, hence facilitating the establishment and maintenance of IBDV persistent infections.

**IMPORTANCE:** Members of the *Birnaviridae* family, including infectious bursal disease virus (IBDV), exhibit a dual behaviour, causing acute infections that are often followed by the establishment of life-long persistent asymptomatic infections. Indeed, persistently infected specimens might act as efficient virus reservoirs, hence potentially contributing to virus dissemination. Despite the key importance of this biological trait, information about mechanisms triggering IBDV persistency is negligible. Our report evidences the capacity of IBDV, a highly relevant avian pathogen, to establishing long-term, productive, persistent infections in both avian and human cell lines. Data presented here provide novel and direct evidence about the crucial role of type I IFNs on the fate of IBDV-infected cells and their contribution to controlling the establishment of IBDV persistent infections. The use of cell lines unable to respond to type I IFNs opens a promising venue to unveiling additional factors contributing to IBDV persistency.

## INTRODUCTION

Viruses are obligate intracellular pathogens establishing complex relationships with their hosts in order to successfully accomplish their main biological roles, i.e. replicating and spreading. Most viruses cause short-lived acute infections ensuing the production of an abundant infective progeny. This enables virus dissemination before the host’s immune system clears the infection. As a general rule, viruses exploiting this strategy are cytocidal. However, under certain circumstances, some lytic viruses, e.g. Reoviruses and Picornaviruses, deviate from this behaviour and establish productive persistent infections in their hosts and/or in cultured cells (1, 2).

Members of the *Birnaviridae* family (3) also exhibit a dual behaviour, causing acute infections that are often followed by the establishment of a life-long persistency. Epitomizing the major biological significance of this trait, birnaviruses belonging to all four family genera have been isolated from asymptomatic carrier specimens including insects (4), fish (5, 6) and birds (reviewed in 7). Despite this, the characterization of molecular basis underlying the initiation and maintenance of birnaviral persistent infections has been customarily overlooked. Hence, current information about this key biological feature, largely shaping birnavirus lifestyles, is as yet rather scarce.

Infectious bursal disease virus (IBDV), the sole member of the *Avibirnavirus* genus, is the best characterized component of the *Birnaviridae* family. IBDV virions are naked icosahedrons (65-70 nm in diameter; T=13L symmetry) harbouring a single capsid enclosing a bipartite double-stranded RNA genome that encodes five mature polypeptides (VP1-VP5) (7).

Although productively infecting a wide variety of bird species, clinical signs of disease have only been reported in juvenile (2-6 weeks old) domestic chickens (*Gallus gallus*). Two IBDV serotypes have been identified this far. Serotype II strains, mainly isolated from asymptomatic turkeys (*Maleagris gallopavo*), are non-pathogenic for chickens. In contrast, all serotype I strains induce clinical signs of disease. However, the virulence of these strains is highly variable, ranging from mild to very virulent. The latter causing very high (nearing 100%) mortality rates in unvaccinated chicken flocks. Indeed, very virulent IBDV strains pose a major threat to the poultry industry world-wide (reviewed in 8).

Several years ago, our group described the establishment of IBDV persistent infections in chicken DT40 cultured cells (9). Regrettably, DT40 cells originated from a chicken lymphoma, hence, *ab initio* being persistently infected with avian leukosis virus (ALV) (10). Indeed, the presence of co-infecting ALV posed a major obstacle to studying the mechanism(s) allowing the establishment of IBDV persistent infections.

Here, we describe an experimental model allowing to generate long-term, stable, persistent IBDV infections in chicken DF-1 and human HeLa cells, both free of endogenous viruses.

Our results show that IBDV infection involves an acute phase characterized by a very active virus replication, the release of high virus titres and a massive apoptotic response that kills most infected cells. Cells surviving the initial acute phase remain persistently infected, sustaining cell proliferation whilst enduring a highly mitigated virus replication. The long-term maintenance of persistently infected DF-1 cells leads to selection of clonal cell populations unable to respond to type I interferon (IFN).

Data collected after the pharmacological elimination of IBDV from persistently infected cells indicate that the functional inactivation of type I IFN signalling drastically increases both the capacity of infected cells to survive the acute IBDV infection phase and their susceptibility to establishing new persistent infections. Work carried out with a knockout human HeLa cell line unable to respond to type I IFN fully recapitulates data gathered with chicken DF-1 cells.

This report provides a novel molecular insight about the crucial role of type I IFNs on the fate of IBDV-infected cells and their contribution to controlling the establishment of IBDV persistent infections.

## RESULTS

### Establishment of IBDV persistent infections

Infection of DF-1 cell monolayers with IBDV invariably leads to a massive cell death within the first 48-72 h post-infection (PI). However, a marginal fraction of infected cells remains alive long after this period. This recurrent observation led us to perform a detailed characterization of this phenomenon.

Pre-confluent DF-1 monolayers were infected at a multiplicity of infection (MOI) of 3 plaque forming units (PFU) per cell with IBDV. Four days later, culture plates were rinsed twice with DMEM to remove cell debris, and then maintained in normal medium. Cell culture medium was routinely replaced every five days. After three weeks, cultures were either stained with crystal violet to visualize surviving cells or processed for immunofluorescence (IF) to assess the expression of the virus-encoded VP3 polypeptide.

As shown in Fig. 1A, after the 21 days “recovery period” culture plates were populated with well differentiated cell islets exhibiting a wide size range. Microscopic analysis indicated that, regardless their size, islets were formed by healthy-looking cells (Fig. 1B) showing a specific VP3 IF signal formed by discrete protein accumulations within the perinuclear region (Fig. 1C). Indeed, VP3 expression indicated that cell islets were formed by persistently infected cells. These cultures are henceforth termed persistently infected DF-1 (DF-1P).

**Figure 1.**
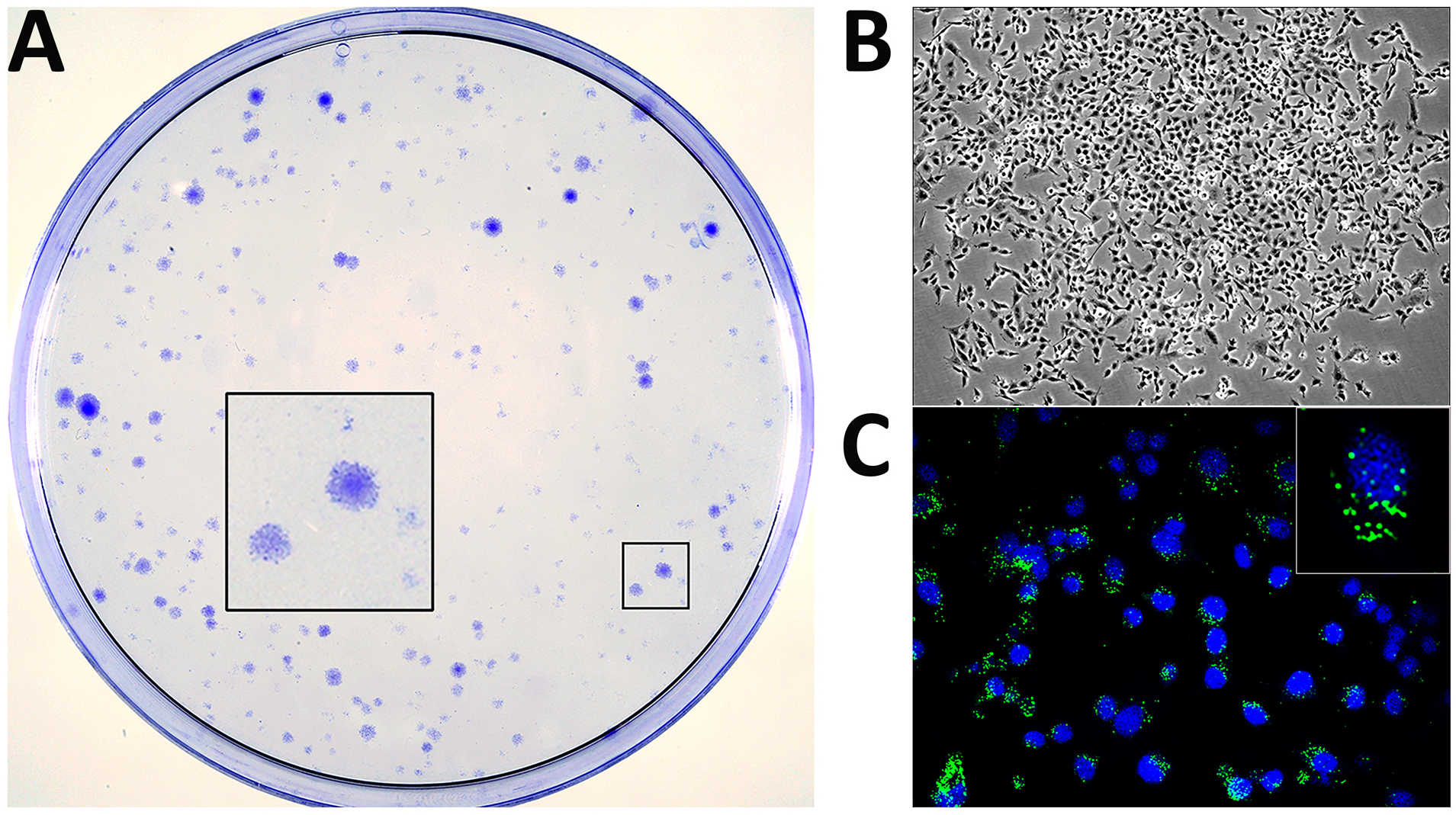
Generation of persistently infected DF-1 cell cultures. DF-1 cell monolayers were infected with IBDV at a MOI of 3 PFU/cell. At four days PI, cultures were carefully rinsed to remove cell debris and then maintained in DMEM supplemented with 10% FCS. Culture media was replaced every five days. At 21 days PI, cultures were either stained with crystal violet to visualize surviving cell clones **(A)** or processed for phase contrast **(B)** or IF microscopy **(C)** after incubation with an antibody specifically recognising the IBDV structural VP3 polypeptide (green). Cell nuclei (blue) were stained with DAPI. Insets show a higher magnification (x2.5) of boxed areas.

Assuming the clonal origin of these cell islets, data gathered from three independent experiments indicate that, under the described experimental conditions, a 0.023±0.007% of infected DF-1 cell population survive the acute IBDV infection phase.

Noteworthy, during the first two-three months, DF-1P cultures undergo sporadic crises resulting in the death of a substantial fraction of the total cell population. However, after this period, cultures become stable maintaining steady proliferation rates.

### Characterization of DF-1P cultures

Three independently generated DF-1P cultures were maintained for a period of 180 days and then used to assess three critical parameters, i.e. expression of virus-encoded proteins, accumulation of viral RNAs and extracellular infectious virus production. Mock- and acutely infected (3 PFU/cell) naïve DF-1 cells collected at 18 h PI were used as controls for these assays.

First, cultures were processed for IF analysis using an anti-VP3 serum. As shown in Fig. 2A, all DF-1P cells showed a positive signal for this polypeptide. Noteworthy, in contrast to the rather intense cytoplasmic immunostaining detected in acutely infected cells, the VP3 signal found in DF-1P cells was characterized by formation of small perinuclear protein accretions. In line with this observation, Western blotting analysis revealed that the accumulation of the VP3 polypeptide in DF-1P cultures was significantly lower than that detected in acutely infected cells (Fig. 2B). We next compared the relative abundance of the IBDV RNA by reverse transcription and quantitative PCR (RT-qPCR) analysis. As shown in Fig. 2C, accumulation of IBDV RNA in DF-1P cultures was ca. three log_10_ units lower than that found in acutely infected cells. Similarly, extracellular IBDV titres detected in DF-1P cell supernatants were ca. four log_10_ units lower that those found in samples from acutely infected DF-1 cells (Fig.2D). Results obtained with the three selected DF-1P cultures were nearly identical.

**Figure 2.**
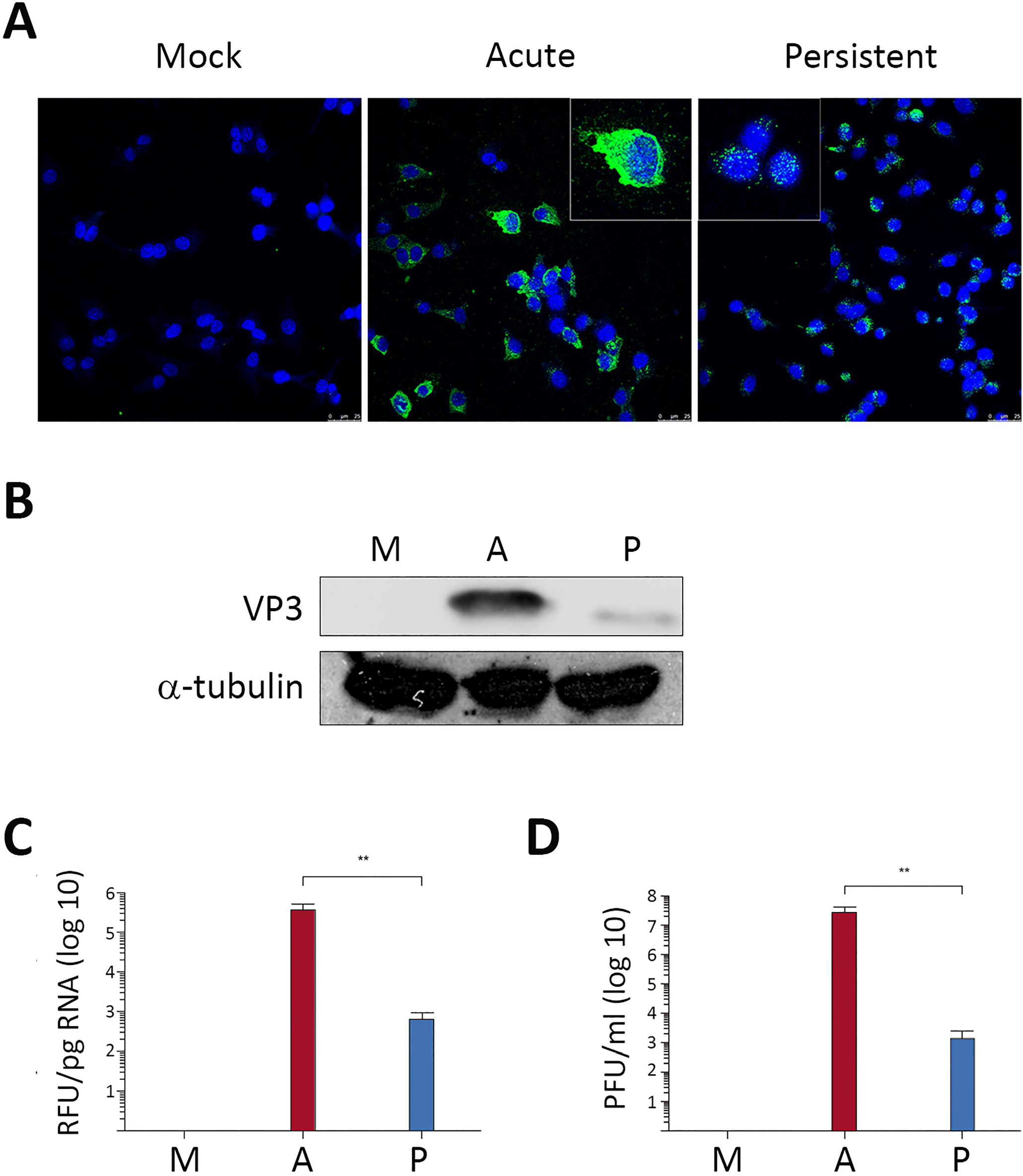
Characterization of persistently infected DF-1P cell cultures. Persistently infected DF-1P cell cultures [P] maintained for 180 days were used to assess the expression of virus-encoded proteins, the accumulation of viral RNAs, and the production of extracellular infectious virus. Mock-[M] and acutely infected (3 PFU/cell) DF-1 cells collected at 18 h PI [A] were used as controls for this analysis. **(A)** Cells were processed for IF microscopy using an antibody specifically recognising the IBDV structural VP3 polypeptide. Cell nuclei (blue) were stained with DAPI. Insets show higher magnification (x2.5) of boxed areas. **(B)** Cell extracts were subjected to SDS-PAGE followed by Western blotting using antibodies specifically recognising either the IBDV structural VP3 or the cellular α-tubulin polypeptides. **(C)** RNAs extracted from the different cell cultures were used for RT-qPCR analysis using primers hybridizing at the VP3 coding region (genome segment A). **(D)** Supernatants collected from the different cell cultures were used for virus titration. Each determination was carried out in triplicate. Presented data correspond to the mean ± the standard deviation of three independent experiments. Brackets indicate pairwise data comparisons. **p<0.001 as determined by two-tailed unpaired Student’s t-test.

### Characterization of viral populations generated during persistent infection

Results described above indicate that both virus replication and infectious virus yields detected in DF-1P cultures are exceedingly lower than those found in acutely infected DF-1 cells. Hence, suggesting the possibility that persistency might entail the selection of viral populations with reduced replication fitness. To analyse this hypothesis, we first determined the nucleotide sequence of the parental (WT) and the DF-1P virus population (P). Then, comparisons of both genome segments and their four virus-encoded polypeptides were performed (Supplemental data file 1). Genome differences were minimal, having identity scores of 97.3 and 98% for segments A and B, respectively. Similarly, rather high identity scores were also detected when comparing for the proteins encoded both genomes; i.e. 97.9, 98.2 and 99.5% for VP5, polyprotein and VP1, respectively. Noteworthy, the C-terminal end of the VP1 polypeptide encoded by the P virus population contains an extra glutamine residue (Q). As far as we know, the presence of C-terminal VP1 extensions has not been reported before. Indeed, it would be worth to determine whether this mutation might affect its RdRp activity.

The growth rates of the WT virus stock and samples collected from supernatants of DF-1P cells (P) were compared. Owing to the rather low infectious titres found in samples from DF-1P cells, this study was performed by infecting naïve DF-1 cell monolayers at a MOI of 0.005 PFU/cell. As shown in Fig. 3, the replication efficiency of the P virus is slightly greater, rendering extracellular infectious titres ca. 1 log_10_ units higher at 48 and 72 h PI, than that detected in cells infected with the parental WT virus. Similar results were obtained with P virus populations harvested from the three independent DF-1P cultures.

**Figure 3.**
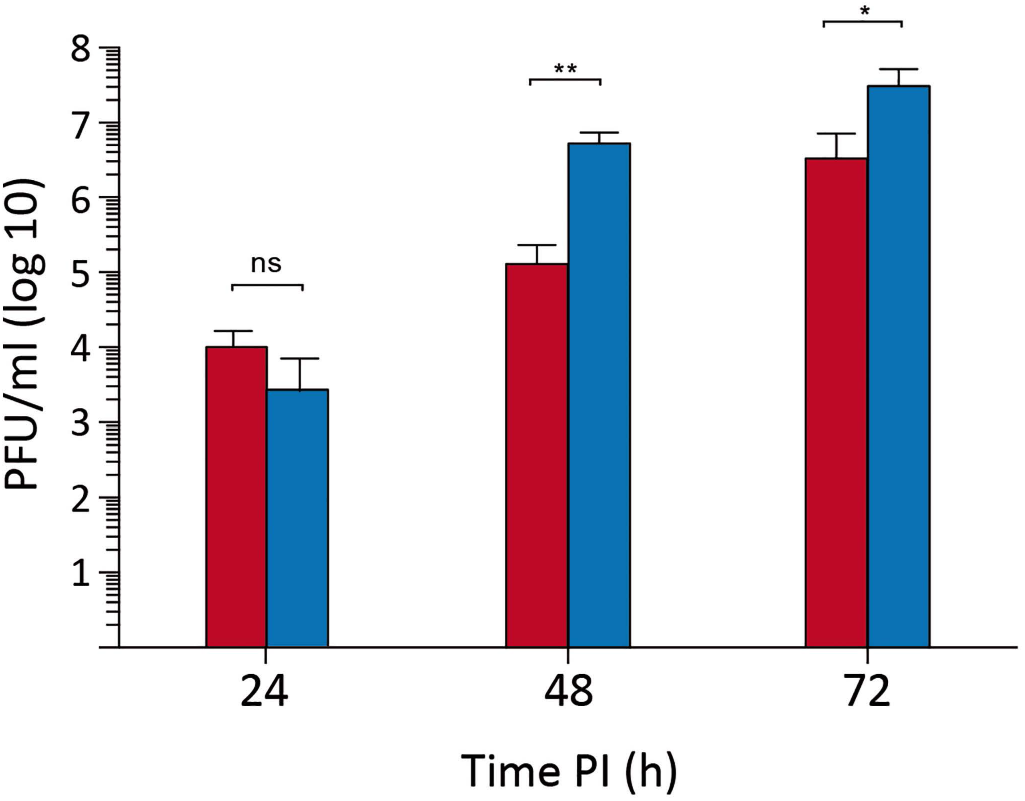
Replication fitness of persistent virus populations. DF-1 cell cultures were infected (0.005 PFU/cell) with the parental IBDV (red) or a supernatant collected from a 180 days old DF-1P culture (blue). Infected cell supernatants were collected at the indicated times PI and used for virus titration. Each determination was carried out in triplicate. Presented data correspond to the mean ± the standard deviation of three independent experiments. Brackets indicate pairwise data comparisons. *p<0.01, **p<0.001 as determined by two-tailed unpaired Student’s t-test. ns, not significant.

Next, we sought to determine whether viral populations collected during persistency exhibited an enhanced capacity to establishing new persistent infections. For this, a series of experiments were carried out by infecting (0.005 PFU/cell) naïve DF-1 cell cultures with either the WT or the P virus. After four days, cultures were rinsed, allowed to recover for three weeks and then used to determine the number of surviving clones as described above. No significant differences were found between WT- or P-infected cells, indicating that both virus populations behave similarly regarding the establishment of persistent infections in DF-1 cells.

### Elimination of infectious IBDV from DF-1P cells

In view of data presented above, it was interesting to compare the behaviour of naïve DF-1 and DF-1P cells. However, the presence of the persistently infecting IBDV population in DF-1P cells posed a major obstacle for this study. Accordingly, we sought to eliminate the virus using 7-Deaza-2’-C-Methyladenosine (7DMA), an efficient inhibitor of a wide variety of viral RNA-dependent RNA polymerases, including that encoded by IBDV (11). DF-1P cultures were maintained for 90 days in medium supplemented with 200 μM 7DMA. Thereafter, cultures were further maintained for 30 days in normal medium to allow the replication of potentially surviving infectious IBDV.

To assess the effect of the 7DMA treatment, cell extracts were used to perform RT-qPCR and Western blotting to search for the presence of IBDV-specific RNA and the VP3 polypeptide, respectively. Samples from untreated DF-1P and acutely-infected (3 PFU/cell) DF-1 cells were used as controls for this analysis. As shown in Fig. 4A and B, the RT-qPCR and Western blotting analyses failed to detect either IBDV-encoded RNA or the VP3 polypeptide in cultures subjected to the described 7DMA treatment. Additionally, infectious IBDV was not found in samples from cell supernatants, indicating that the treatment had effectively eliminated IBDV from DF-1P cultures. Cured DF-1P cultures are hereafter termed DF-1PC.

**Figure 4.**
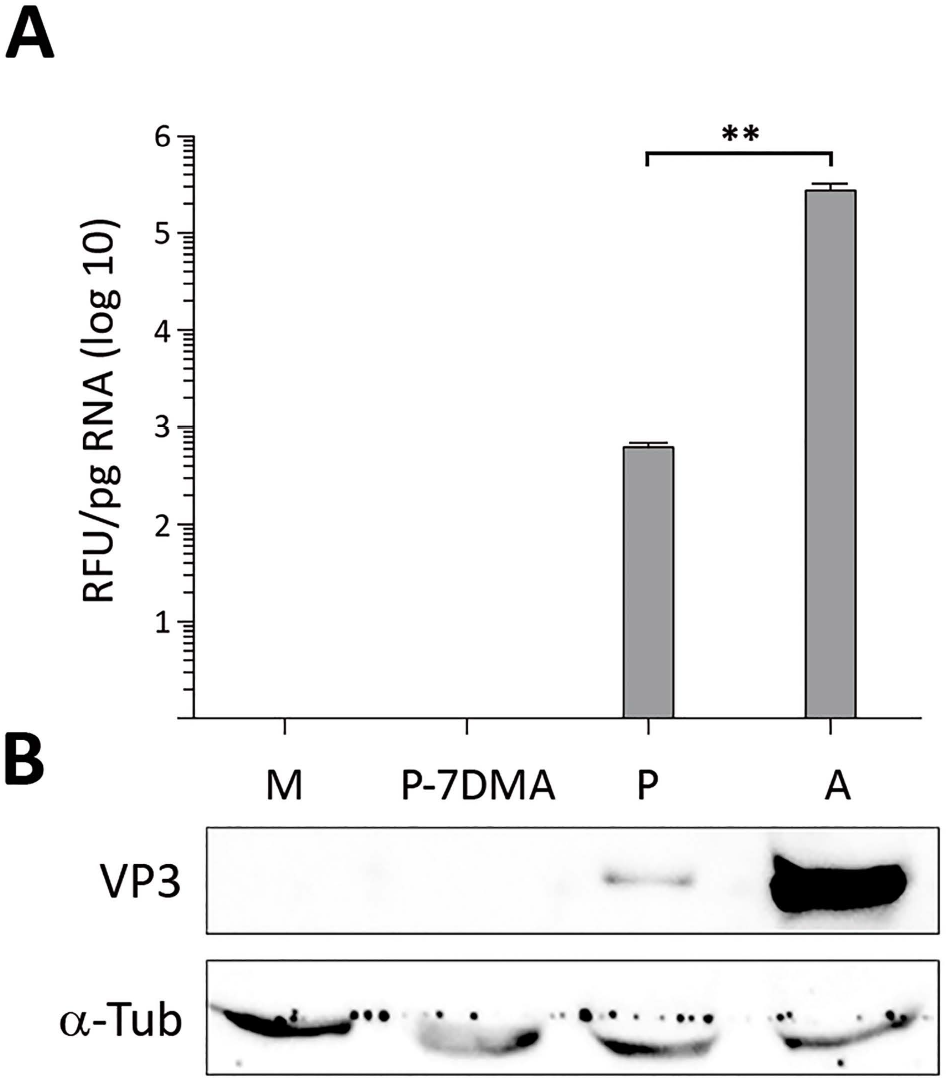
Elimination of infectious IBDV from DF-1P cells. Persistently infected DF-1P cultures were subjected to a prolonged (180 days) treatment with 7-DMA, an inhibitor of viral RNA polymerases of RNA viruses. After the treatment, cultures were maintained for 30 days in the absence of the inhibitor to facilitate the recovery of potentially surviving IBDV populations. Thereafter, untreated [P] and 7DMA-treated [P-7DMA] DF-1P Cell cultures were used to search for the presence of IBDV RNA and the structural IBDV VP3 polypeptide. Samples from mock-[M] and acutely infected DF-1 [A] cells were used as controls. **(A)** RNAs extracted from the different cultures were used to perform a RT-qPCR analysis using primers hybridizing at the VP3 coding region (genome segment A). Each determination was carried out in triplicate. Presented data correspond to the mean ± the standard deviation of three independent experiments. Brackets indicate pairwise data comparisons. **p<0.001 as determined by two-way ANOVA. **(B)** Cell extracts were subjected to SDS-PAGE followed by Western blotting using antibodies specifically recognising either the IBDV structural VP3 or the cellular α-tubulin polypeptides.

### DF-1PC cells exhibit an enhanced susceptibility to establishing persistent IBDV infections

At this point, it was interesting to comparing the growth rate of WT IBDV in naïve DF-1 and DF-1PC cell cultures. Accordingly, both cell lines were infected with WT IBDV at a MOI of 3 PFU/cell. Cell supernatants were collected at different times (i.e. 24, 48 and 72 h) PI and used for virus titration. As shown in Fig. 5, no significant differences were observed on the extracellular infectious yields generated upon infection of these two cell lines.

**Figure 5.**
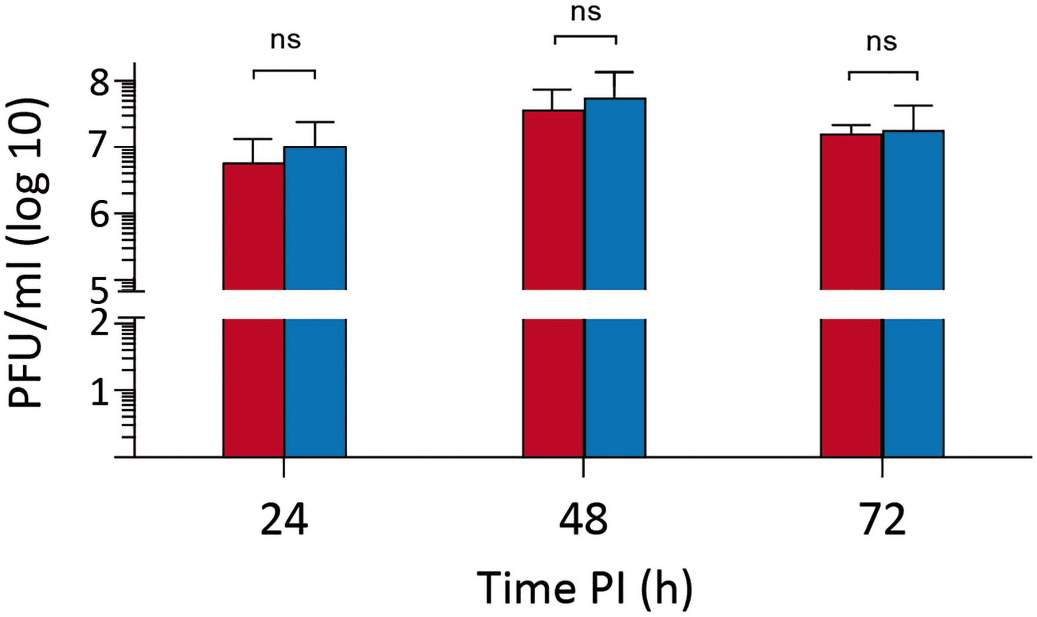
Susceptibility of IBDV-cured DF-1PC cells to IBDV infection. Naïve DF-1 (blue) and DF-1PC (red) cultures were infected with WT IBDV (3 PFU/cell). Cell supernatants were harvested at the indicated times PI and used for virus titration. Presented data correspond to the mean ± the standard deviation of three independent experiments. Brackets indicate pairwise data comparisons. **p<0.001 as determined by two-way ANOVA. ns, not significant.

Despite the efficient IBDV replication observed with DF-1PC cells, it was readily noticeable that the cell death induced by the infection was much lower in DF-1PC than in the parental DF-1 cell line. To quantitatively assessing this initial observation, a kinetic cell death analysis was performed using the MTT assay. DF-1 and DF-1PC cell monolayers were infected with WT IBDV (3 PFU/cell), and cell viability determined at 1, 3 and 6 days PI, respectively. As shown in Fig. 6A, infection of parental DF-1 cells led to a swift reduction of cell viability, reaching undetectable MTT values at 3 days PI. In contrast, the viability decline of infected DF-1PC cells was significantly less pronounced, reaching its minimal value (ca. 35%) at 3 days PI. Furthermore, MTT values detected in infected DF-1PC cultures underwent a substantial increase at 6 days PI, indicating that cells surviving the acute infection phase maintained the capacity to proliferate. Fig. 6B shows representative phase contrast images corresponding to IBDV-infected DF-1 and DF-1PC cultures captured at 1 h and 6 days PI, respectively, evidencing the existence of a major difference on the capacity of these two cell lines to survive an acute IBDV infection phase. Noteworthy, infected DF-1PC cultures re-established fully confluent persistently infected monolayers 8-10 days after the initiation of the infection.

**Figure 6.**
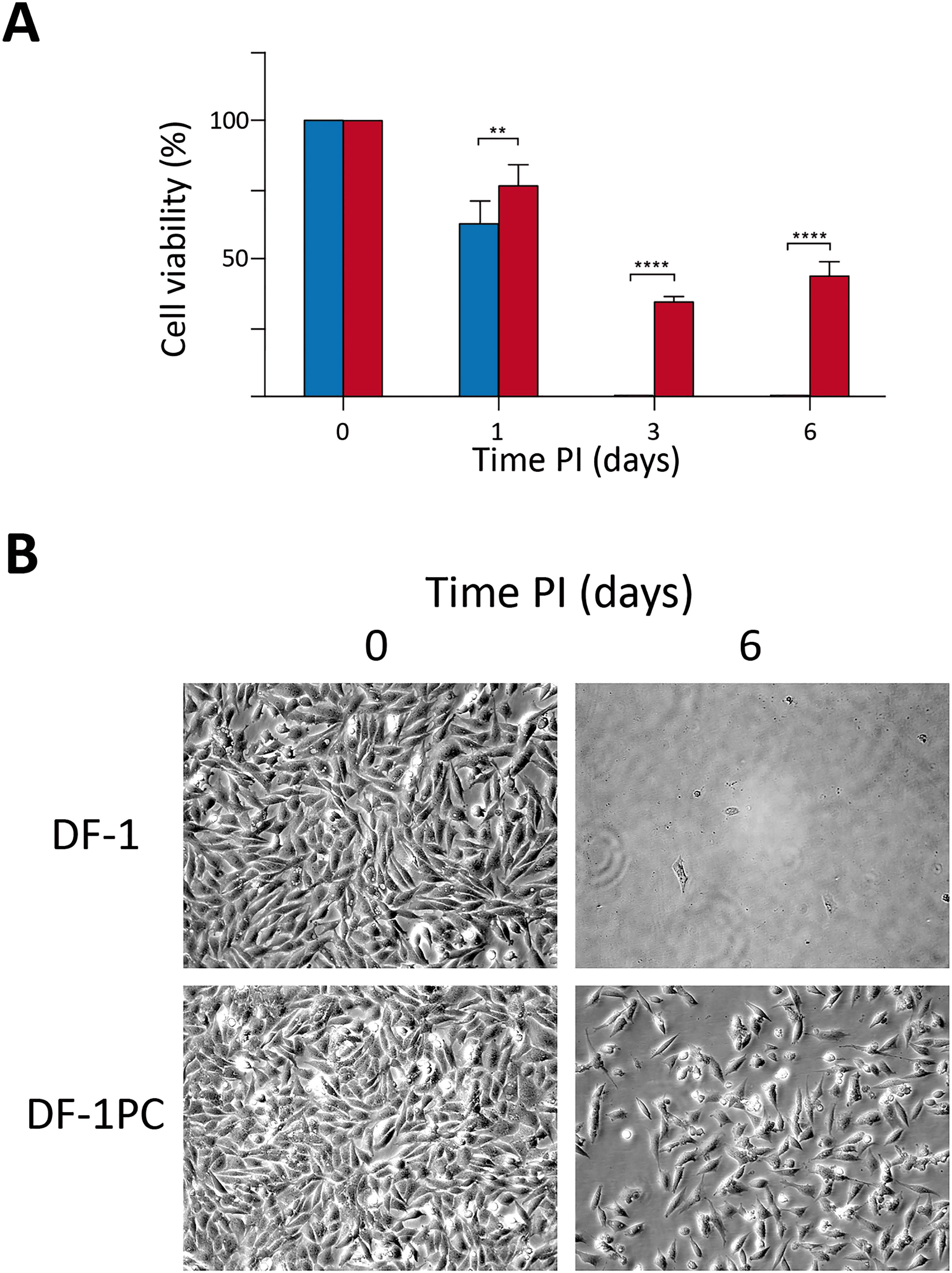
Comparative analysis of the capacity of naïve DF-1 and DF-1PC cells to survive the acute IBDV infection phase. DF-1 (blue) and DF-1PC (red) cell cultures were infected with WT IBDV (3 PFU/cell). **(A)** Cell viability was determined at the indicated times PI using the MTT assay. Presented data correspond to the mean ± the standard deviation of three independent experiments. Brackets indicate pairwise data comparisons. **p<0.001, ****p<0.00001 as determined by two-way ANOVA. **(B)** Phase contrast microscopy images of infected DF-1 and DF-1PC cell cultures captured at the indicated times PI.

### The JAK-STAT signalling pathway is functionally blocked in DF-1PC cells

We have recently shown that type I IFN plays a crucial role in the fate of IBDV-infected cells (11). Thus, whilst pre-treatment of cells with IFNα induces a solid protection against IBDV replication, its addition early after infection leads to a massive apoptotic response that wipes out infected cell cultures (11).

It was therefore important to compare the capacity of DF-1 and DF-1PC cells to respond to type I IFN. To this end, DF-1 and DF-1PC cultures were treated or not with IFNα (1,000 IU/ml) during 16 h. Thereafter, cultures were incubated with a recombinant vesicular stomatitis virus expressing the GFP protein (VSV-GFP) (1 PFU/cell). VSV is extremely sensitive to the antiviral response induced by type I IFNs, hence, providing a simple assay to assessing IFN responsiveness. At 12 h PI, cultures were incubated with DAPI to stain cell nuclei and then visualized using fluorescence microscopy to detect GFP expression (Fig. 7A). As expected, untreated DF-1 cells showed an intense GFP signal, thus indicating that they were efficiently infected with VSV-GFP. In contrast, IFNα-treated DF-1 cells were fully protected, as evidenced by the absence of a fluorescent GFP signal. Interestingly, both untreated and IFNα-treated DF-1PC showed an intense green fluorescence revealing the VSV-GPF infection, hence indicating the incapacity of this cell line to respond to IFNα.

**Figure 7.**
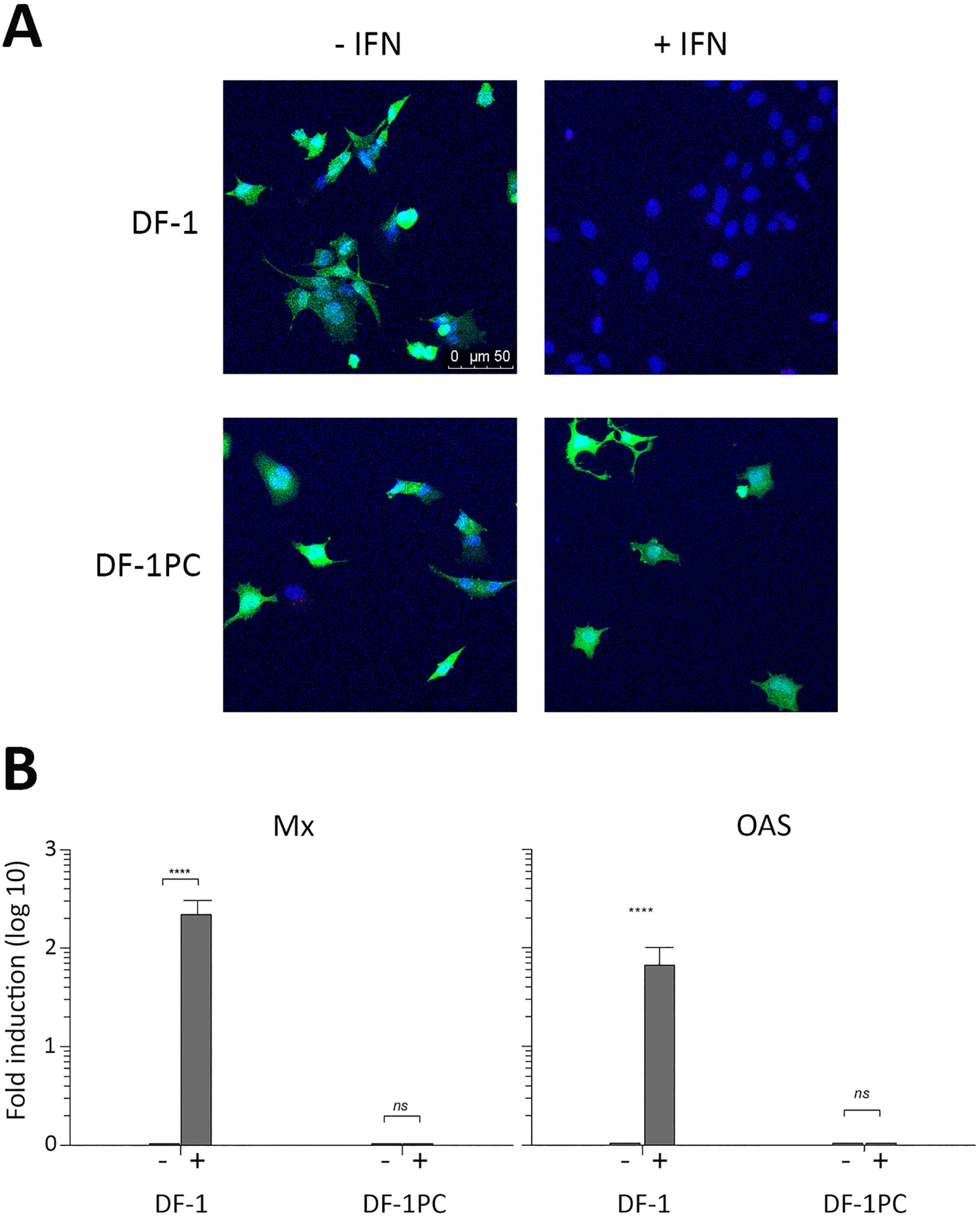
Determination of the capacity of naïve DF-1 and DF-1PC cells to respond to chicken IFNα. DF-1 and DF-1PC cell cultures were treated (*IFN) or not (-IFN) with of IFNα (1,000 IU/ml) during 16 h. Cultures were then used to assess their susceptibility to VSV infection or the expression of the Mx and OAS ISGs. **(A)** Treated and untreated cultures were infected (1 PFU/cell) with VSV-GFP expressing the GFP polypeptide. At 12 h PI, cultures were incubated with DAPI to stain cell nuclei and then observed under fluorescent microscopy. **(B)** Treated and untreated cultures were harvested and subjected to RNA extraction. Mx and OAS expression levels were determined by RT-qPCR. Presented data correspond to the mean ± the standard deviation of three independent experiments. Brackets indicate pairwise data comparisons. ****p<0.00001 as determined by two-way ANOVA. ns, not significant.

To further verify this finding, an alternative approach was used. DF-1 and DF-1PC monolayers were treated or not with IFNα (1,000 IU/ml) for 16 h, and then used to extract cellular RNA. These RNA samples were used to perform RT-qPCR analyses to check the transcriptional activation of two interferon-stimulated genes (ISG), i.e. Mx and 2’,5’-oligoadenylate synthetase (OAS), generally used as activation markers of the Janus kinase-signal transducer and activator of transcription protein (JAK-STAT) signalling pathway (12). As shown in Fig. 7B, DF-1 cells efficiently responded to the IFNα treatment triggering a strong transcriptional activation of both Mx and OAS genes. In contrast, basal Mx and OAS RNA levels were detected in both treated and untreated DF-1PC cells. These results further confirmed the incapacity of the DF-1PC cell line to respond to IFNα.

### Genetic analysis of DF-1 and DF-1PC cells

The avian JAK-STAT signalling pathway is relatively simple, encompassing a reduced number of protein components (reviewed in 13). Despite this apparent simplicity, there are many potential mutations capable of impairing the pathway’s functionality. In view of this, high-throughput sequencing (NGS) was used to perform a whole sequencing analysis of the DF-1 and DF-1PC cell genomes as described in the Material and Methods section.

Assembled DF-1 and DF-1PC genomes were annealed to the *Gallus gallus* (breed Red Jungle Fowl isolate RJF #256 [accession GRCg6a]) genome. Both assembled genomes cover over 98% of the reference genome with an average sequencing depth of 35x (Supplementary Table 1). Next, DF-1 and DF-1PC exomes were compared to the reference chicken exome to detect differential single nucleotide polymorphisms (SNPs), insertions and deletions. This analysis led to the detection of two specific SNPs exclusively found within the DF-1PC exome. Both SNPs consisted in a thymidine (T) per guanine (G) substitution at nucleotide positions 106,590,841 (SNP1) and 106,591,114 (SNP2), respectively. As summarized in Table 1, SNP1 and 2 were found within 57% and 46%, respectively, of DF-1PC sequencing reads spanning the described nucleotides.

**Table 1.**
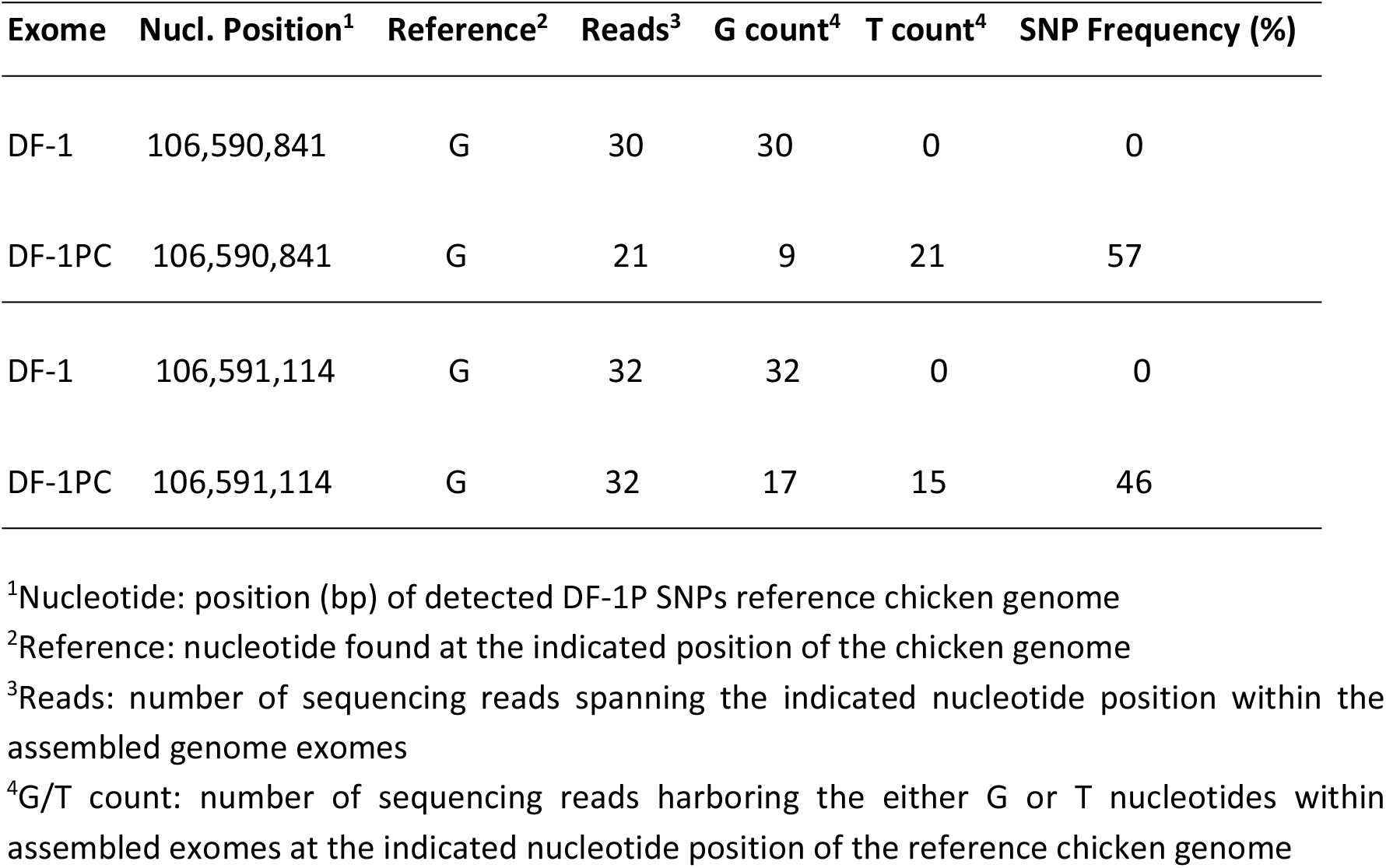
Exome DF-1P SNPs.

| Exome | Nucl. Position<sup>1</sup> | Reference<sup>2</sup> | Reads<sup>3</sup> | G count<sup>4</sup> | T count<sup>4</sup> | SNP Frequency (%) |
| --- | --- | --- | --- | --- | --- | --- |
| DF-1 | 106,590,841 | G | 30 | 30 | 0 | 0 |
| DF-1PC | 106,590,841 | G | 21 | 9 | 21 | 57 |
| DF-1 | 106,591,114 | G | 32 | 32 | 0 | 0 |
| DF-1PC | 106,591,114 | G | 32 | 17 | 15 | 46 |

<sup>1</sup>Nucleotide: position (bp) of detected DF-1P SNPs reference chicken genome
<sup>2</sup>Reference: nucleotide found at the indicated position of the chicken genome
<sup>3</sup>Reads: number of sequencing reads spanning the indicated nucleotide position within the assembled genome exomes
<sup>4</sup>G/T count: number of sequencing reads harboring the either G or T nucleotides within assembled exomes at the indicated nucleotide position of the reference chicken genome

As depicted in Scheme 1, both DF-1PC-specific mutations map at chromosome 1 (NCBI Reference Sequence: NC_006088.05) within the last exon (exon 10) of the gene encoding the IFNα/β receptor subunit 2 (IFNAR2) (GenBank: AF082665.1) (14).

In order to confirm NGS sequencing data, amplicons spanning the IFNAR2 exon 10 region holding both mutations were generated by PCR using genomic DNA isolated either from DF-1 and DF-1PC cells as templates. Oligonucleotides OL_IFNAR2.1 (5’-CCATCCCATCAGCCTGGAAAT) and OL_IFNAR2.2 (5’-TGCACATTGCCAGTCAACAG), hybridizing at nucleotide positions 660-688 and 1349-1368, respectively, of the IFNAR2 DNA were used as PCR primers. DF-1- and DF-1PC-derived amplicons were subjected to conventional nucleotide sequencing analysis using oligonucleotides OL_IFNAR2.1 and OL_IFNAR2.2. As shown in Fig. 8, the presence of double nucleotide (G/T) signals at SNP1 and 2 positions in chromatograms from DF-1PC amplicons fully confirmed NGS data. As expected, amplicons derived from DF-1 genomic DNA presented single (G) nucleotide signals at equivalent positions.

**Figure 8.**
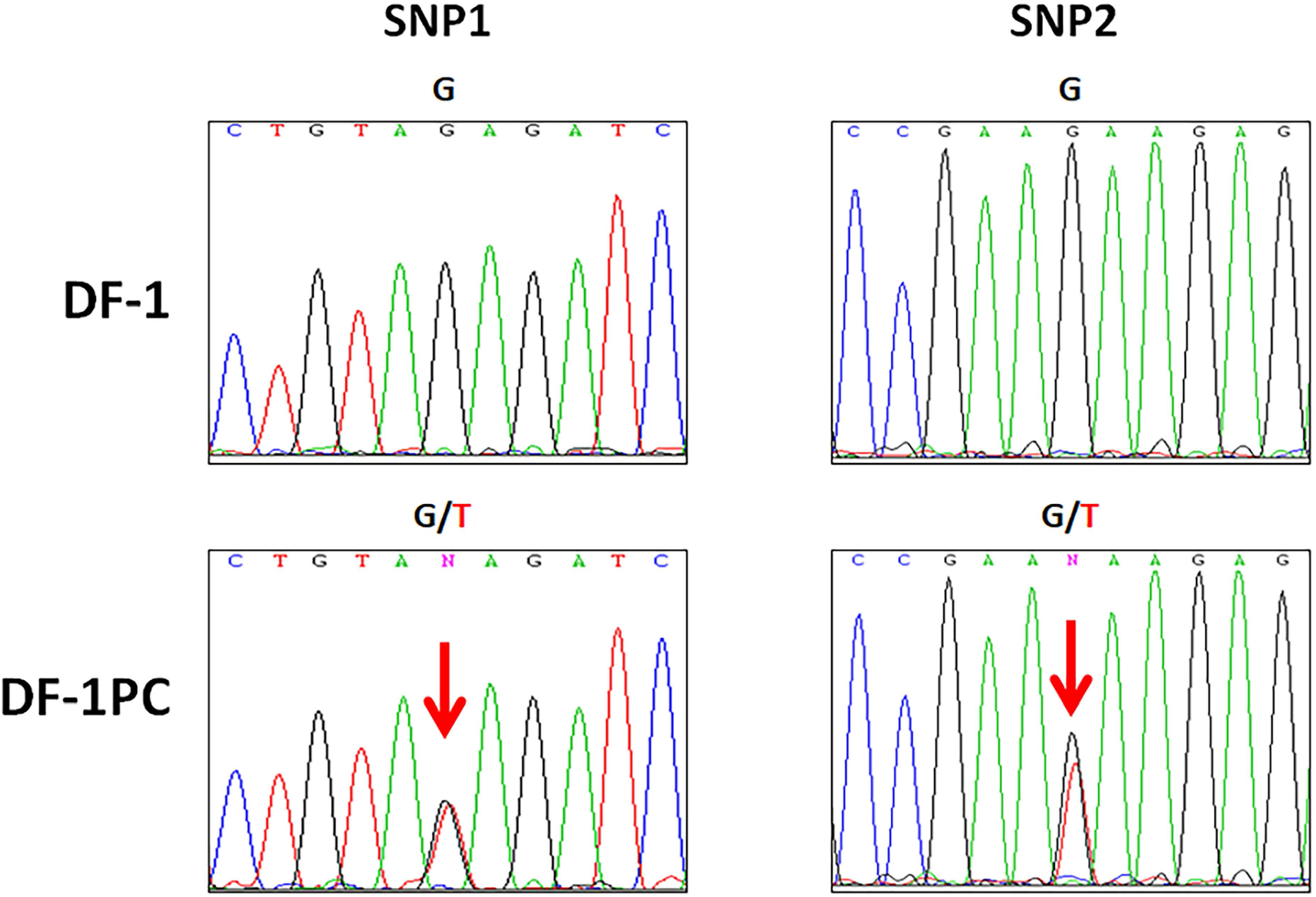
Detection of single nucleotide polymorphisms in IFNAR exon 10. Genomic DNA isolated from DF-1 and DF-1PC cell cultures were subjected to PCR using oligonucleotides OL_IFNAR2.1 and OL_IFNAR2.2. The resulting amplicons, spanning the interest region of IFNAR2 exon 10, were subjected to conventional Sanger sequencing using the same oligonucleotides. Panels show sequencing chromatogram regions of interest (holding DF-1PC SNPs) from DF-1 and DF-1PC amplicons, respectively. Lettering above chromatograms correspond to detected nucleotides, namely A (green), C (blue), G (black) and T (red). Sequence ambiguities (G/T) are indicated as N (magenta).

At this point, it was important to determine whether the detected mutations were homo- or heterozygous. To answer this question, the DF-1PC amplicon was cloned into the pGEM-T Easy plasmid vector. Thereafter, recombinant plasmids purified from 20 independent ampicillin-resistant *E. coli* colonies were purified and used for nucleotide sequencing using the above described primers. All analysed plasmids contained single mutations; 9 of them harbouring SNP1, and the remaining 11 SNP2. These results indicate that mutations are heterozygous, each DF-1PC IFNAR2 allele containing a single G/T substitution at different exon 10 positions.

The three persistently infected DF-1P cell lines were shown to harbour the same mutations, thus indicating that both mutations were already present before the 7-DMA treatment.

### DF-1PC cells encode C-terminal truncated IFNAR2 polypeptides

The chicken IFNAR2 gene encodes the IFNAR2 protein precursor, a polypeptide (508 residues) encompassing a signal peptide which is removed during its transport to the cell membrane. The mature IFNAR2 is a type I transmembrane polypeptide including a N-terminal ectodomain (216 residues), a transmembrane region (20 residues) and a cytoplasmic domain (246 residues). The ectodomain is responsible for the interaction of the protein with its cognate cytokine ligands and the subsequent assembly of ternary receptor complexes with IFNAR1. IFNAR1 and IFNAR2 cytoplasmic domains are accountable for the propagation of the activation signal to effector proteins, i.e. JAK, STAT1 and TYK2 (reviewed in 15).

The G per T nucleotide substitutions found in the IFNAR2 of DF-1P and DF-1PC cells transform GAT and GAA codons, encoding two glutamic acid [E] residues, i.e. E301 and E392, into TAG and TAA translation termination codons, respectively. Indeed, these mutant IFNAR2 genes encode C-terminal truncated IFNAR2 polypeptides lacking either 107 and 208 residues, respectively (Scheme 2). Unfortunately, the lack of specific antibodies recognizing the chicken IFNAR2 polypeptide prevented to visualize the truncated proteins.

The ablation of the IFNAR2 cytoplasmic domain, essential for the transduction of the JAK-STAT activation signal (16), would explain the incapacity of DF-1PC cells to respond to type I IFN. To confirm this prediction, we sought to determine whether the ectopic expression of a full-length version of the IFNAR2 polypeptide may rescue IFNα responsiveness. Hence, a recombinant version of the chicken IFNAR2 gene was synthesized and cloned into the expression vector pCI-neo, thus generating the pCI-neo/chIFNAR2-Flag plasmid. DF-1PC cells were transfected either with empty pCI-neo or pCI-neo/chIFNAR2-FLAG. At 24 h post-transfection, cultures were treated or not with 1,000 IU of IFNα for 16 h and then used for RNA extraction. RNA samples were used to assess Mx and OAS transcription by RT-qPCR. RNAs from treated and untreated DF-1 cell cultures were used as controls. As shown in Fig. 9, whilst pCI-neo-transfected DF-1PC cultures did not respond to the treatment, those transfected with pCI-neo/chIFNAR2-FLAG readily reacted to IFNα, triggering a ca. 100xfold increase on the expression of both ISGs. These results indicate that the ectopic expression of the full-length recombinant IFNAR2-Flag protein restores the functionality of the JAK-STAT pathway in DF-1PC cells. Additionally, these results ruled out the existence of potentially undetected genetic alterations also affecting the functionality of this pathway.

**Figure 9.**
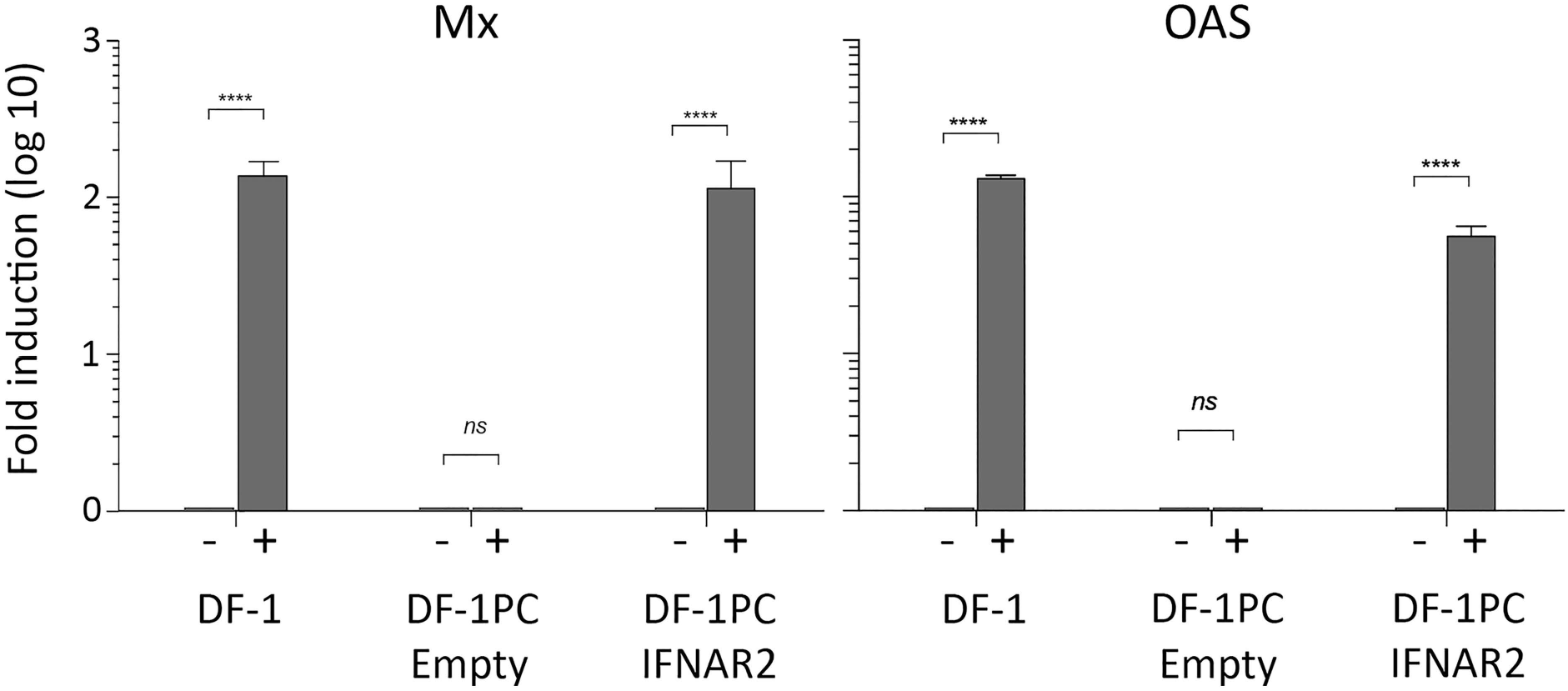
Rescuing DF-1PC JAK-STAT pathway activity. DF-1PC cells were transfected with pCI-neo/chIFNAR2-FLAG, a mammalian expression vector expressing a full length version of the chicken IFNAR2 gene (DF-1PC IFNAR2). Control DF-1PC cultures were transfected with the pCI-neo parental vector (DF-1PC Empty). At 24 h post-transfection, cultures were treated (+) or not (−) with 1,000 IU of IFNα for 16 h. IFN-treated and untreated DF-1 cell cultures were used as additional controls for these experiments. RNA isolated from the different cell cultures was used to assess the transcriptional activation of the Mx and OAS genes by RT-qPCR. Presented data correspond to the mean ± the standard deviation of three independent experiments. Brackets indicate pairwise data comparisons. ****p<0.00001 as determined by two-way ANOVA. ns, not significant.

### Effect of the impairment of the JAK-STAT pathway on the establishment of IBDV persistent infections on human HeLa cells

We have shown that the inactivation of the JAK-STAT pathway significantly lessens IBDV-induced cell death in avian DF-1 cells and enhances their proneness to initiate persistent infections. It was then important to ascertain whether or not this is a general trait also operating in other IBDV-susceptible cells. Concerning this, Urin et al. (17) have recently described the generation of a set of human HeLa knockout cell lines selectively lacking every gene participating in the JAK-STAT pathway. We borrowed the HeLa IFNAR2 KO line, unable to respond to type I IFN, to analyse the effect that the functional inactivation of the JAK-STAT pathway may have on the fate of IBDV-infected HeLa cells.

We first compared the death rate of parental (WT) and IFNAR2 KO cells induced by the IBDV infection using the MTT cell viability assay. Cell monolayers were infected (3 PFU/cell), and cell viability was determined at 1, 3 and 6 days PI. As shown in Fig. 10A, the behaviour of both cell lines was strikingly different. WT cells were swiftly killed by the infection, reaching viability values nearing 0% at 3 days PI. In contrast, the viability of infected IFNAR2 KO HeLa cells remained largely unaltered at the beginning of the infection (1-3 days PI) and grew at later times, hence reflecting the capacity of infected cells to proliferate. In line with this observation, infected IFNAR2 KO cells readily became persistently infected. As it is the case with DF-1P cells, persistently infected IFNAR2 KO HeLa cells also showed a reduced accumulation of the IBDV VP3 polypeptide when compared to acutely infected cells (Fig. 10B).

**Figure 10.**
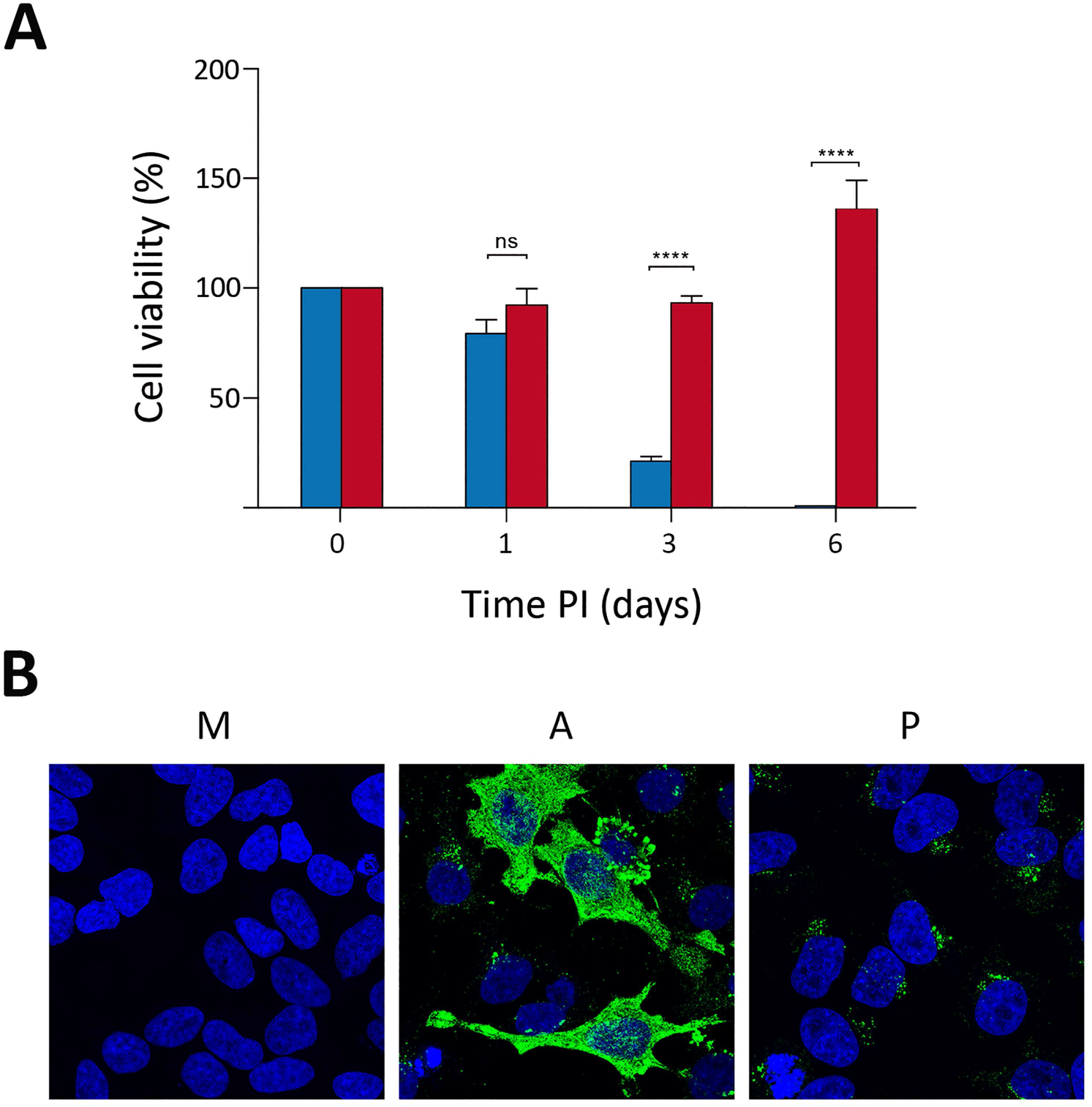
Effect of the inactivation of the JAK-STAT pathway on the fate of IBDV-infected HeLa cells. **(A)** WT (blue) and IFNAR2 KO HeLa (red) cell cultures were infected with WT IBDV (3 PFU/cell). Cell viability was determined at the indicated times PI using the MTT assay. MTT values recorded immediately after infection were considered as 100% cell viability. Presented data correspond to the mean ± the standard deviation of three independent experiments. Brackets indicate pairwise data comparisons. ****p<0.00001 as determined by two-way ANOVA. ns, not significant. **(B)** Persistently infected IFNAR2 KO cells [P] maintained for two months were processed for IF analysis using an antibody specifically recognising the IBDV structural VP3 polypeptide. Cell nuclei (blue) were stained with DAPI. Mock-infected [M] and acutely infected (3 PFU/cell, fixed at 48 h PI) [A] cells were used as controls.

To further assess the differential behaviour of WT and IFNAR2 KO HeLa cells, a Western blotting analysis was performed using cell extracts collected at different times PI (i.e. 24, 48 and 72 h). Samples from mock-infected cells were used as controls. As shown in Fig. 11, both WT and IFNAR2 KO HeLa cells were readily infected with IBDV, thus showing similar VP3 accumulation levels at 24 h PI. As expected, while infection of WT cells efficiently triggered a robust Mx protein expression, infected IFNAR2 KO cell extracts were void of this protein. Thus, confirming their incapacity to activate the type I IFN-mediated gene expression in response to IBDV infection. The proteolytic cleavage of Poly(ADP-ribose) polymerase (PARP) is generally used as an early apoptotic marker (19). Accordingly, we monitored PARP cleavage to comparing the apoptotic response of infected WT and IFNAR2 KO cells. As expected, infection of WT cells resulted in a conspicuous PARP cleavage, detectable from 24 h PI onwards. In contrast, this effect was significantly abridged in IFNAR2 KO cells, showing a rather modest increase of the cleaved PARP (c-PARP) product at 72 h PI. Consistent with the observed apoptotic induction observed in WT HeLa cells, the amount of both VP3 and the cellular actin polypeptide, used as a protein loading control in this experiment, was significantly reduced in WT-infected cells collected at 72 h PI.

**Figure 11.**
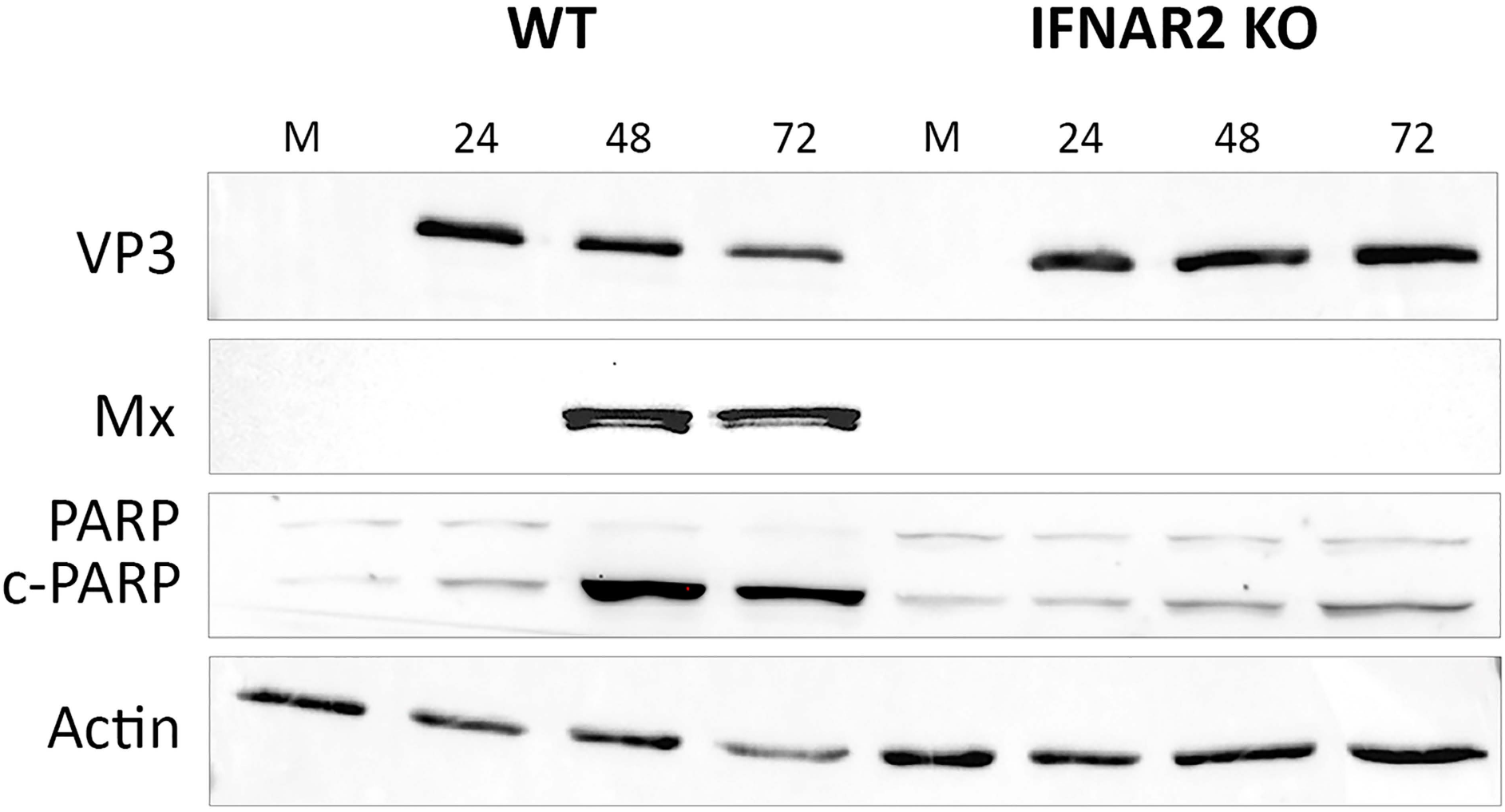
The apoptotic HeLa cell response induced by IBDV infection is largely dependent upon the activation of the JAK-STAT pathway. Infected (3 PFU/cell) WT and IFNAR2 KO cell cultures were collected at the indicated times PI. Mock-infected [M] cell cultures were used as controls. The corresponding extracts were subjected to SDS-PAGE followed by Western blotting using antibodies against the virus-encoded VP3 and the cellular Mx, PARP and actin proteins. The PARP cleavage product is denoted as c-PARP.

Taken together, these results show that, similarly to what had been found in DF-1 cells, the obliteration of the JAK-STAT signalling pathway significantly enhances the capacity of HeLa cells to survive the acute infection phase and facilitates the establishment of IBDV persistent infections.

## DISCUSSION

Although infection of DF-1 cells with IBDV results in a massive cell death, a minor, yet consistent, cell fraction endures the acute infection phase giving rise to a collection of persistently infected cell clones, holding from ten to several thousand cells at 21 days PI. This initial size variation suggests the existence of major differences in the capacity of surviving cells to withstand IBDV replication. Our results indicate that continuous cell passaging progressively narrows down the initial heterogeneity. Hence, although persistently infected cultures described here were not subjected to cell cloning, genome sequencing data showed that the three DF-1P lines characterized during our study share identical mutations on the IFNAR2 gene, suggesting that they originated from a pre-exiting DF-1 cell subset already holding these mutations.

It seems reasonable to hypothesize that the long-term passaging of persistently infected cultures enables a Darwinian process resulting in the selection of clonal cell populations harbouring the fittest genotype to sustain proliferation whilst enduring a significantly lessened, yet productive, IBDV replication. Indeed, the repeated selection of persistently infected cell clones incapable to respond to type I IFN described here underscores the crucial role of the innate antiviral cell response and, in particular that of the JAK-STAT pathway, on the fate of IBDV-infected cells.

### Virus replication taming, an IBDV persistency hallmark

As evidenced in different virus-cell systems, persistent viral infections require the successful interplay of cellular and viral mechanisms capable of sustaining a precise equilibrium between virus replication and cell proliferation. The modulation of virus replication is a common finding in persistent infections, likely being a necessary requirement for the initiation of persistency (19).

A comparison of elemental virus replication parameters (i.e. virus yield, genome and protein accumulation) performed in acute and persistent IBDV infections indicates that persistency entails a major downregulation of virus replication. IBDV downregulation could have been associated to the selection during the initial stages of the persistency of a virus population(s) exhibiting a reduced replication fitness. However, contrary to this simple notion, viruses harvested from persistently infected DF-1 cells showed an enhanced replication capacity when compared to the WT parental virus. A similar situation was found during the characterization of BHK-21 cells persistently infected with foot-and-mouth disease virus (FMDV; *Picornaviridae*) (20). In both cases, IBDV and FMDV, persistency results in the selection of virus populations exhibiting an enhanced fitness.

Although the possible implication of other genetic elements, e.g. microRNAs (21), cannot be ruled out at this point, data gathered from a wide variety of viruses, including birnaviruses, suggest that the downregulation of IBDV replication might be related to the presence of defective virus genomes (DVGs). Although these aberrant genomes, arising from errors during the replication of many viruses, harbor protein coding defects, DVGs retain the replication capacity of standard viral genomes. It has been extensively documented that DVGs interfere the replication of parental viruses, hence contributing to the establishment of persistent infections both *ex* and *in vivo* (22, 23). Moreover, it has been recently shown that DVGs from respiratory syncytial virus (*Pneumoviridae*) and Sindbis virus (*Alphaviridae*), are key players in a complex mechanism that fools the innate antiviral cell response, specifically promoting the survival of persistently infected cells (24). Indeed, this observation constitutes a major conceptual leap further highlighting the DVGs’ relevance on virus-host interactions and on their central role in shaping viral lifestyles.

Although a molecular analysis was missing, an early report from Müller et al. (25), suggested the accumulation of DVGs and defective interfering virus particles during IBDV replication. Additionally, several reports concerning the establishment of IPNV, a closely related birnavirus, persistent infections also advocated the possible role of DVGs on this phenomenon (26, 27). Unfortunately, the characterization of birnavirus DVGs has been discontinued for over 30 years. Indeed, data described in this report has revived our interest for this phenomenon, hence experiments aimed to ascertain the possible contribution of DVGs to the establishment of persistent IBDV infections are currently ongoing.

### The role of the JAK-STAT pathway

The type I IFN-dependent antiviral response entails two interrelated signalling cascades known as transactivation and JAK-STAT pathways, respectively. The transactivation pathway, involving a large number of sentinel and effector proteins, is activated by the presence of pathogen associated molecular patterns (e.g. nucleic acids, proteins or lipids) at different cell compartments, triggering the expression of type I IFNs as well as an ISG subset, including IFN response factors, pattern recognition receptors and some antiviral effectors (13). Although for the sake of concision, data concerning the transactivation capacity of DF1-PC and HeLa are not presented in this report, we have documented that this pathway remains fully functional in both DF-1PC and HeLa IFNAR2 KO cell lines.

Upon their release from infected cells IFNs interact with both uninfected (paracrine IFN activity) and infected (autocrine IFN activity) cells. This interaction activates the JAK-STAT pathway, triggering the expression of the whole ISG set and the concomitant implementation of the antiviral defence programme (Schneider et al., 2014). The autocrine/paracrine feed-back allows type I IFNs to generate an antiviral state in uninfected bystander cells. Most relevant for our study, the autocrine IFN activity further boosts the transactivation pathway on infected cells, hence initiating the so-called autocrine signalling loop (28). It has been well documented that the concurrent activation of both type I IFN pathways induces a vigorous apoptotic response that eliminates virus-infected cells (reviewed in 29). Indeed, the comparative analysis performed with DF-1 and DF-1PC cells indicates that the functional inactivation of the JAK-STAT pathway significantly reduces the apoptotic response induced by the infection strongly affecting IBDV-induced death rates. Thus, while MTT cell viability values recorded in WT infected DF-1 cells become negligible at 72 h PI, approximately 30% of infected DF-1PC cells remain alive at this time. A comparison of data corresponding to the recovery of persistently infected cell clones generated upon infection of naïve DF-1 cells, a 0.023% of the total infected population, with the 30% cell survival recorded in infected DF-1PC populations, indicates that the inactivation of the JAK-STAT pathway enhances over 1,000-fold the capacity of DF-1 cells to survive the acute IBDV infection phase.

Results obtained with WT and IFNAR2 KO HeLa cells nicely recapitulate the differential behaviour detected in infected DF-1 and DF-1PC cells, showing again that the inactivation of the JAK-STAT pathway affords a striking increase on the capacity of infected cells to survive the acute infection phase and to initiate persistent infections. In line with previous reports, results presented here show that the death of IBDV-infected cells is directly related to the apoptotic response triggered by the infection, and that this is largely owed to the autocrine activity of type I IFNs. Indeed, a major conclusion of our study is that type I IFNs act as a major barrier preventing the establishment of IBDV persistent infections.

As stated above, IBDV replication in HeLa is significantly slower than in DF-1 cells. A major difference is also evident in the production of infectious virus yields, being significantly higher (ca. 1.5-2 log_10_ units) in DF-1 cells. Interestingly, despite the fact that both cell lines, DF1-PC and HeLa IFNAR2 KO, share the incapacity to respond to type I IFN, the efficiency with which HeLa IFNAR2 KO develop persistent infections is significantly higher than that observed in DF-1PC cells. This observation suggests the possible existence of an inverse correlation between replication efficiency and the proneness to establishing persistent infections.

Although IgM-bearing chicken bursal lymphocytes are the main IBDV cell target, it has been shown that, yet with a much lower efficiency, the virus also replicates in cells of the monocyte-macrophage lineage (30). It is tempting to speculate with the possibility that this cell lineage could play a role in the establishment of *in vivo* persistent infections.

### Additional factors contributing to the apoptotic response in IBDV-infected cells

Our data indicates that the functional disruption of the JAK-STAT pathway does not completely preclude IBDV-mediated apoptosis. Hence, suggesting that other factor(s) associated to IBDV replication process also contribute to this phenomenon.

It can be envisaged that the hefty accumulation of both virus dsRNA and proteins, and the assembly of huge cytoplasmic arrays holding thousands of closely packed virions, along with the appropriation of the biosynthetic machinery during the acute IBDV infection phase can significantly alter cellular homeostasis, thus affording additional proapoptotic stimuli. Regarding this, the availability of IFNα/β-unresponsive DF-1PC and HeLa IFNAR2 KO cells might help in the identification of other factors, viral and/or cellular, contributing to the obliteration of IBDV-infected cells. Indeed, both cell lines provide a powerful tool to further dissect the molecular bases of the IBDV-induced cellular pathogenesis.

### Concluding remarks

As recently evidenced with Ebola virus (EBOV), persistently-infected asymptomatic individuals support the long-term maintenance and resurgence of EBOV epidemics in apparently virus-free geographical areas (31). Regarding IBDV, the relentless reemergence of virus outbreaks, even in areas under intense vaccination programs and strict hygienic measures, suggests the existence of undetected virus reservoirs. Indeed, persistently infected birds might play an important role on IBDV epidemiology. Hopefully, data presented here might help in stimulating the interest on this largely overlooked phenomenon.

## MATERIAL AND METHODS

### Cells, viruses and infections

DF-1 (chicken embryonic fibroblasts, ATCC number CRL-12203) and HeLa (human epithelial cervical cancer cells) were grown in Dulbecco’s modified Eagle’s medium (DMEM) supplemented with penicillin (100 U/ml), streptomycin (100 mg/ml) and 5% fetal calf serum (FCS) (Sigma). IBDV infections were performed on preconfluent (ca. 75%) cell monolayers with the Soroa strain, a cell-adapted serotype I virus, diluted in DMEM at an MOI of 3 PFU per cell, unless otherwise stated. After adsorption (1 h, 37°C), the medium was replaced with fresh DMEM supplemented with 2% FCS. Infected cultures were maintained at 37°C. Infections with the recombinant stomatitis vesicular stomatitis virus expressing the GFP protein (VSV-GFP) (32) were performed in the same way.

### Virus titrations

For IBDV titrations, supernatants from cultures infected with IBDV were collected and subjected to centrifugation (5,000xg for 5 min) at 4°C to remove cell debris. Clarified cell supernatants were used to determine extracellular virus titers by plaque assay using semisolid agar overlays followed by immunostaining as previously described (33).

### IFNα

The recombinant chicken IFN-α used in our experiments was expressed, purified and titrated in our laboratory as previously described (11).

### Light microscopy

Cell cultures grown onto glass coverslips were washed with PBS, fixed with 4% paraformaldehyde (Sigma) for 30 min and then extensively rinsed with PBS. Cells were permeabilized by incubation with PBS containing Triton X-100 (Sigma) 0.5% for 5 min. Coverslips were blocked for 20 min using a solution of PBS containing 5% FCS and then incubated with a rabbit anti-VP3 serum (34) for 18 h at 4°C. Thereafter, coverslips were repeatedly washed in PBS, and incubated with goat anti-rabbit IgG coupled to Alexa-488 diluted in PBS supplemented with 1% FCS for 45 min at 20°C. Cell nuclei were stained with 2-(4-Amidinophenyl)-6-indolecarbamidine dihydrochloride (DAPI; Sigma) diluted in PBS for 30 min at 20°C. Finally, coverslips were dehydrated with ethanol and mounted with ProLong antifade reagent (Invitrogen). Cells infected with VSV-GFP were washed with PBS, fixed with 4% paraformaldehyde (Sigma) for 30 min and, after an extensive wash with PBS, cell nuclei were stained as described above. Samples were visualized by epifluorescence using a Leica TCS-Sp5 microscope confocal system. Fluorescent signals detected by CLSM were recorded separately by using appropriate filters. Images were captured using the LASAF v.2.6.0 software package (Leica Microsystems).

### Western blotting

Samples used for Western blot analyses were prepared by removing media from cell monolayers, and then suspending cells in ice-chilled disruption buffer (0.5% Triton X-100; 50 mM KCl; 50 mM NaCl; 20 mM Tris-HCl [pH 7.5]; 1 mM EDTA, 10% glycerol; complete protease inhibitor cocktail [Roche]). Samples were mixed (v/v) with 2x Laemmli’s sample buffer and heated at 95°C for 5 min before electrophoresis. Electrophoreses were performed on 12% polyacrylamide gels. Gels were subjected to electroblotting onto Hybond-C nitrocellulose membranes. Membranes were blocked with 5% nonfat dry milk in PBS for 1 h at room temperature and then incubated 18 h at 5°C with the appropriate antisera. Thereafter, membranes were extensively washed with PBS and incubated with the corresponding secondary antibodies coupled to horseradish peroxidase. Immunoreactive bands were detected by chemiluminescence (Amersham ECL Prime Western Blotting Detection Reagent). Antibodies used in our experiments are listed in Table 2.

**Table 2.**
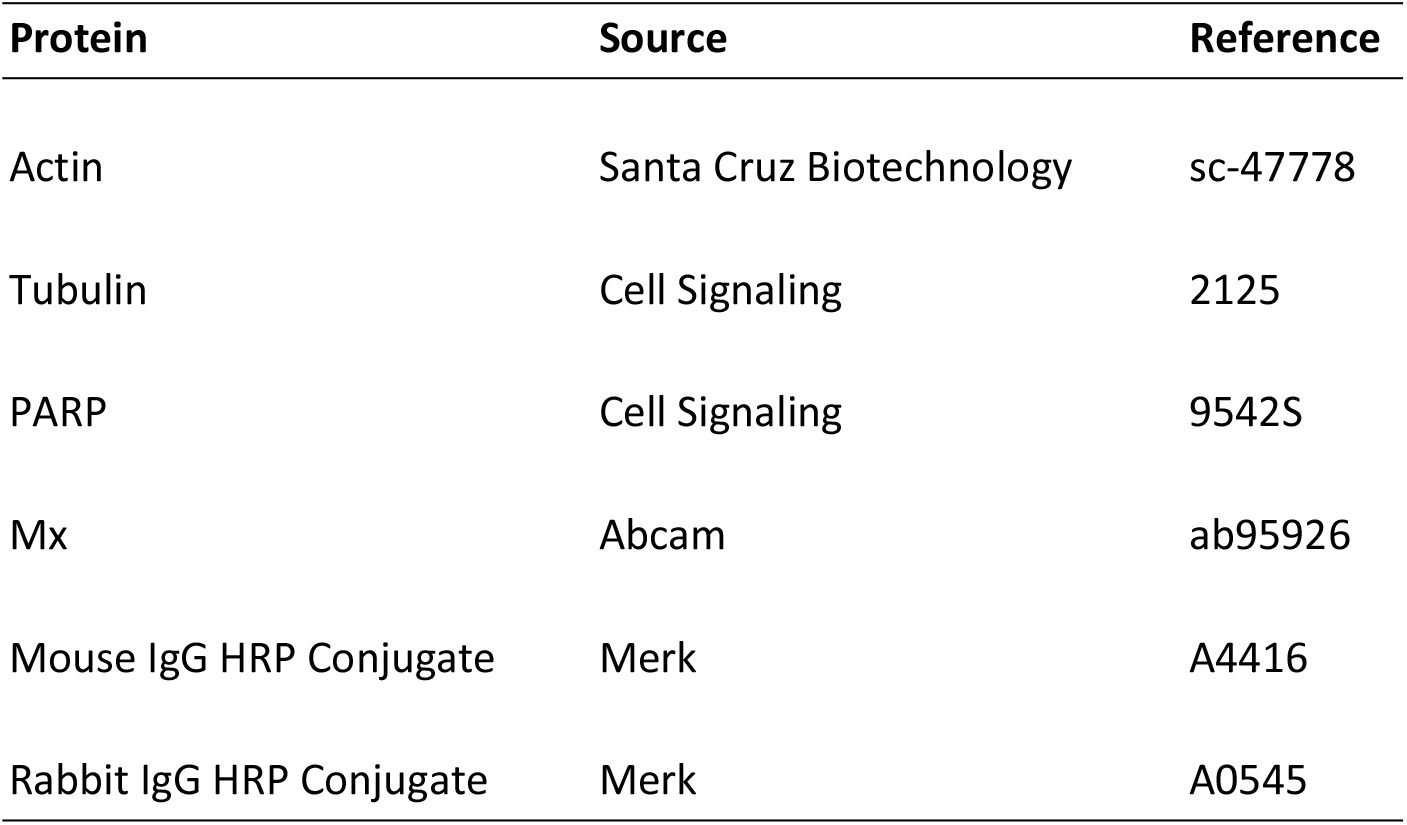
List of antibodies used for Western blotting analysis.

### Quantitative RT-qPCR analysis

Total RNA was isolated by using the Macherey-Nagel™ NucleoSpin RNA Plus XS kit (ThermoFisher Scientific) according to the manufacturer’s instructions. Purified RNAs (300-500 ng) were reverse transcribed into cDNA by using SuperScript III (Invitrogen) reverse transcriptase and random primers. The resulting cDNA samples were subjected to qPCR using the gene-specific primers listed in Table 3. Reactions were performed in triplicate by using Power SYBR green PCR master mix (ThermoFisher Scientific), according to the manufacturer’s protocol, and by using an Applied Biosystems 7500 real-time PCR system instrument. Reactions were performed as follows: 2 min at 50°C; 10 min at 95°C; 40 cycles of 15 s at 95°C and 1 min at 60°C; and, finally, 15 s at 95°C, 1 min at 60°C, 30 s at 95°C, and 15 s at 60°C to build the melt curve. Gene expression levels were normalized to the hypoxanthine phosphoribosyl transferase 1 (HPRT) gene, and the results were calculated as fold changes in gene expression relative to mock-infected cells by using the delta-delta CT (threshold cycle) method of analysis.

**Table 3.**
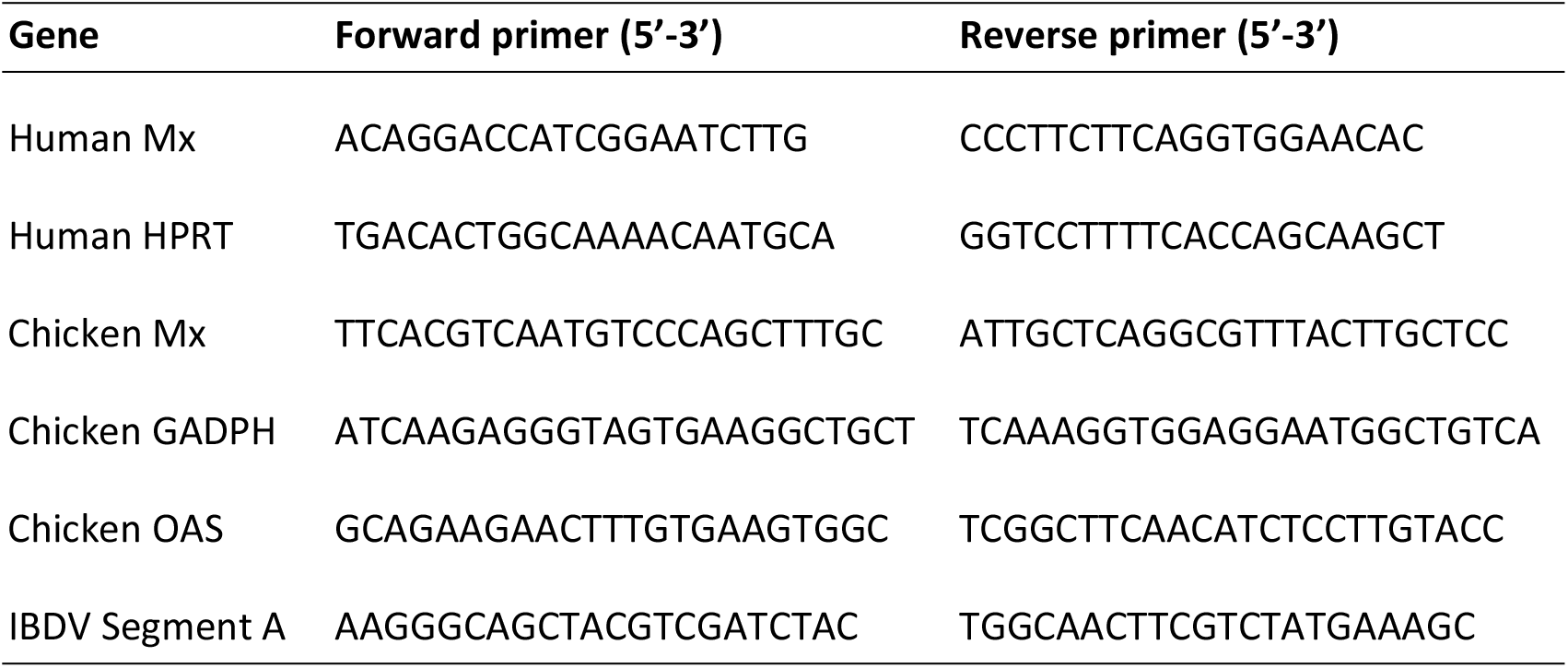
List of primers used for RT-qPCR.

### Cell viability assays

Cell viability was determined using the MTT Assay Kit (Abcam), based on the conversion of water soluble 3-(4,5-dimethylthiazol-2-yl)-2,5-diphenyltetrazolium bromide (MTT) to an insoluble formazan product, following the manufacturer’s instructions. Cell monolayers were incubated for 3 h with DMEM supplemented with the MTT reagent. After this period, cell medium was replaced by plain DMEM and further incubated for 30 min. Thereafter, media were replaced by dimethyl sulfoxide (DMSO) and maintained for 15 min. All incubations were carried out in the dark under normal culture conditions (37°C, 5% CO_2_). Finally, samples were harvested and used to determine the absorbance (Abs) at 570 nm. The percentage of viable cells was determined using the following equation: % viable cells=(Abs_sample_-Abs_blank_/Abs_control_-Abs_blank_)x100. Controls correspond to cell cultures collected at the beginning of the experiment. Blank correspond to empty culture wells.

### IBDV genome sequencing

Complete genome sequencing of viruses was performed as previously described (35) with minor modifications. Briefly, viral RNA was extracted from either DF-1 cultures infected with the WT virus or DF-1P cells using the Qiamp viral RNA mini kit (Qiagen) according to the manufacturer’s instructions. Each segment genome segment was RT-PCR amplified in two fragments using previously described primers (35). Full-length sequencing of each segment was performed in both directions on three independently produced PCR products.

### High-throughput cell genome analysis

Genomic DNA was isolated from DF-1 and DF-1PC cell cultures using the Wizard Genomic DNA Purification kit following the manufacturer’s instructions. A total amount of 1.5]μg DNA per sample was used as input material for the DNA sample preparations. Sequencing libraries were generated using TruSeq Library Construction Kit (Illumina) following manufacturer’s recommendations and index codes were added to attribute sequences to each sample. Briefly, the DNA samples were fragmented by sonication to a size of 350 bp, end-polished, A-tailed, and ligated with the full-length adaptor for Illumina sequencing with further PCR amplification. After the library was constructed, a preliminary quantification was performed using Qubit 2.0 (ThermoFisher Scientific), and the library concentration was diluted to 1 ng/μl. Then, libraries were analyzed for size distribution using an Agilent2100 Bioanalyzer and quantified using real-time PCR. After the library was qualified, library preparations were sequenced on an Illumina HiSeq platform and paired-end reads were generated. The original data obtained from the high throughput sequencing were transformed to sequenced reads by base calling. Raw data were recorded in a FASTQ file containing sequenced reads and corresponding sequencing quality information. The sequenced reads were filtered to remove low quality reads and adapters. After this process, clean reads were mapped to the reference genome for subsequent variation analysis. The bwa software (36) was used for the comparison of short reads obtained from high-throughput sequencing to the *Gallus gallus* (breed Red Jungle Fowl isolate RJF #256 [accession GRCg6a]) genome. The position of clean reads on the reference genome was determined by alignment, and the information such as the depth of sequencing of the samples and the genome coverage were counted and used for variation detection. A summary of the statistics of the alignments with Reference Genome is shown in Supplemental Table 1. Individual SNP and InDels were detected using SAMTOOLS (37) and annotated using the ANNOVAR software (38). Library construction, high-throughput sequencing, sequence alignments and SNP and InDel detection and annotations were performed by CD Genomics (NY, USA).

### PCR cloning and sequencing

Genomic DNA fragments generated by PCR were cloned into the pGEM-T Easy plasmid vector using the pGEM®-T Easy Vector System (Promega) following the manufacturer’s instructions. Cloned DNAs were subjected to nucleotide sequencing using the SP6 and T7 sequencing primers.

## AKNOWLEDGEMENTS

We are grateful to the excellent technical assistance provided by Antonio Varas, and work of Silvia Gutiérrez-Erlandsson and Ana M. Oña from the Advanced Light Microscopy scientific CNB service. We are also extremely grateful to Prof. Gideon Schreiber for generously sharing WT and IFNAR2 KO HeLa cell lines.

---

**Scheme 1.**
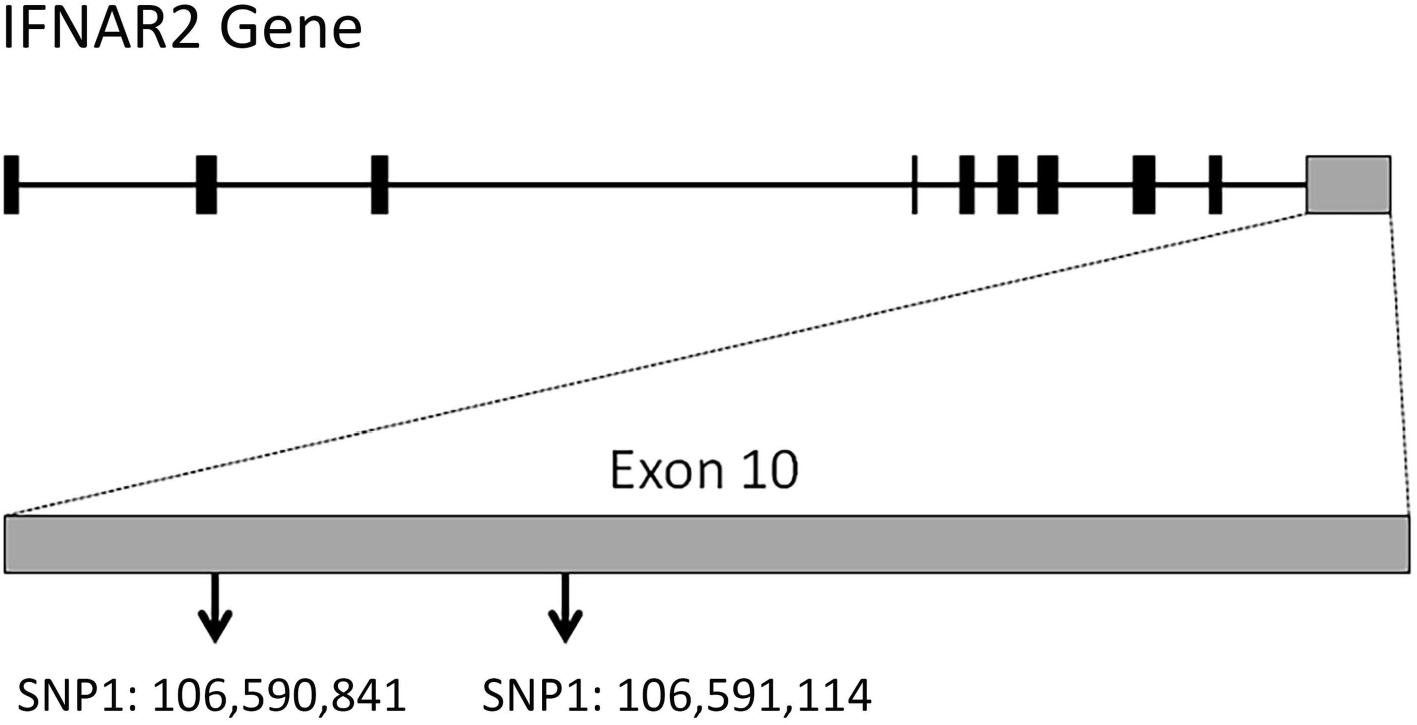
Mapping of DF-1PC IFNAR2 mutations. Cartoon depicting the structure of the chicken IFNAR2 gene. The gene, located at chicken chromosome 1, holds ten exons (indicated as boxes). DF-1PC IFNAR2 mutations detected by NGS map at exon 10 (grey box) at the indicated nucleotides of the reference *Gallus gallus* (breed Red Jungle Fowl isolate RJF #256 [accession GRCg6a]) reference genome.

**Scheme 2.**
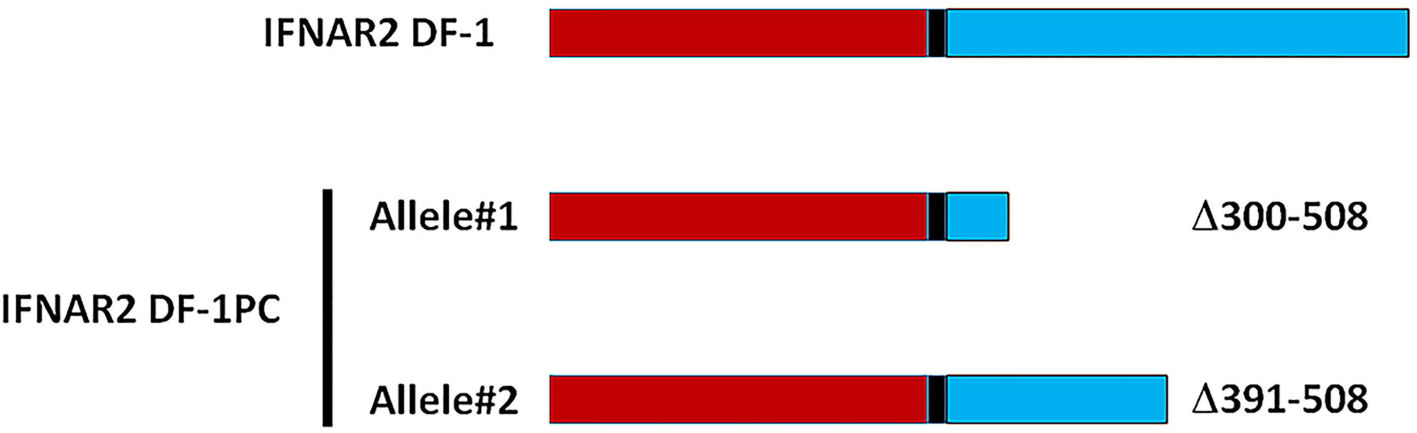
DF-1PC IFNAR2 mutant polypeptides. Cartoon depicting mature IFNAR2 polypeptides expressed by DF-1 and DF-1PC cells, respectively. Noteworthy, DF-1PC IFNAR2 gene alleles encode C-terminal truncated proteins lacking either 107 or 208 residues. The mature IFNAR2 extracellular, transmembrane and cytoplasmic domains are labelled in red, black and blue, respectively.

