## Supplemental Table for "Type I Interferon acts as a major barrier to the establishment of infectious bursal disease virus (IBDV) persistent infections"

| Sample | Total reads (bp) | Mapped reads (bp) | Mapping rate (%) | Average depth (X) | Coverage at least 1X (%) | Coverage at least 4X (%) |
|---|---|---|---|---|---|---|
| DF-1 | 267,004,156 | 263,082,947 | 98.53 | 32.02 | 98.39 | 97.67 |
| DF-1PC | 303,910,872 | 299,145,387 | 98.43 | 35.75 | 98.42 | 97.72 |

**Supplemental Table 1.** Summary of the statistics of alignment results. DF-1 and DF1-PC sequencing reads were mapped to the *Gallus gallus* genome.
