## Supplemental Data for "Type I Interferon acts as a major barrier to the establishment of infectious bursal disease virus (IBDV) persistent infections"

**Supplemental File 1**

### 1. Virus nucleotide sequences

#### 1.1. IBDV WT Segment A

GGATACGATCGGTCTGACCCGGGGGAGTCACCCGGGGACAGGCTGACAAGGCCTTGTTCCAGGATGGAA
CTCCTCCTTCTACAACGCTATCATTGATGGTTAGTAGAGATCAGACAAACGATCGCAGCGATGACAAACC
TGCAAGATCAAACCCAACAGATTGTTCCGTTCATACGGAGCCTTCTGATGCCAACAACCGGACCGGCGTC
CATTCCGGACGACACCCTGGAGAAGCACACTCTCAGGTCAGAGACCTCGACCTACAATTTGACTGTGGGG
GACACAGGGTCAGGGCTAATTGTCTTTTTCCCTGGATTCCCTGGCTCAATTGTGGGTGCTCACTACACAC
TGCAGGGCAATGGGAACTACAAGTTCGATCAGATGCTCCTGACTGCCCAGAACCTACCGGCCAGTTACAA
CTACTGCAGGCTAGTGAGTCGGAGTCTCACAGTGAGGTCAAGCACACTTCCTGGTGGCGTTTATGCACTA
AACGGCACCATAAACGCCGTGACCTTCCAAGGAAGCCTGAGTGAACTGACAGATGTTAGCTACAATGGGT
TGATGTCTGCAACAGCCAACATCAACGACAAAATTGGGAACGTCCTAGTAGGGGAAGGGGTCACCGTCCT
CAGCTTACCCACATCATATGATCTTGGGTATGTGAGGCTTGGTGACCCCATTCCCGCAATAGGGCTTGAC
CCAAAAATGGTAGCCACATGTGACAGCAGTGACAGGCCCAGAGTCTACACCATAACTGCAGCCGATGATT
ACCAATTCTCATCACAGTACCAACCAGGTGGGGTAACAATCACACTGTTCTCAGCCAACATTGATGCCAT
CACAAGCCTCAGCGTTGGGGGAGAGCTCGTGTTTCGAACAAGCGTCCACGGCCTTGTACTGGGCGCCACC
ATCTACCTCATAGGCTTTGATGGGACAACGGTAATCACCAGGGCTGTGGCCGCAAACAATGGGCTGACGA
CCGGCACCGACAACCTTATGCCATTCAATCTTGTGATTCCAACAAACGAGATAACCCAGCCAATCACATC
CATCAAACTGGAGATAGTGACCTCCAAAAGTGGTGGTCAGGCAGGGGATCAGATGTCATGGTCGGCAAGA
GGGAGCCTAGCAGTGACGATCCATGGTGGCAACTATCCAGGGGCCCTCCGTCCCGTCACGCTAGTGGCCT
ACGAAAGAGTGGCAACAGGATCCGTCGTTACGGTCGCTGGGGTGAGCAACTTCGAGCTGATCCCAAATCC
TGAACTAGCAAAGAACCTGGTTACAGAATACGGCCGATTTGACCCAGGAGCCATGAACTACACAAAATTG
ATACTGAGTGAGAGGGACCGTCTTGGCATCAAGACCGTCTGGCCAACAAGGGAGTACACTGACTTTCGTG
AATACTTCATGGAGGTGGCCGACCTCAACTCTCCCCTGAAGATTGCAGGAGCATTCGGCTTCAAAGACAT
AATCCGGGCCATAAGGAGGATAGCTGTGCCGGTGGTCTCCACATTGTTCCCACCTGCCGCTCCCCTAGCC
CATGCAATTGGGGAAGGTGTAGACTACCTGCTGGGCGATGAGGACCAGGCCGCTTCAGGAACTGCTCGAG
CCGCGTCAGGAAAAGCAAGAGCTGCCTCAGGCCGCATAAGGCAGCTGACTCTCGCCGCCGACAAGGGGTA
CGAGGTAGTCGCGAATCTATTCCAGGTGCCCCAGAATCCCGTAGTCGACGGGATTCTTGCTTCACCTGGG
GTACTCCGCGGTGCACACAACCTCGACTGCGTGTTAAGAGAGGGTGCCACGCTATTCCCTGTGGTTATTA
CGACAGTGGAAGACGCCATGACACCCAAAGCATTGAACAGCAAAATGTTTGCTGTCATTGAAGGCGTGCG
AGAAGACCTCCAACCTCCATCTCAAAGAGGATCCTTCATACGAACTCTCTCTGGACACAGAGTCTATGGA
TATGCTCCAGATGGGGTACTTCCACTGGAGACTGGGAGAGACTACACCGTTGTCCCAATAGATGATGTCT
GGGACGACAGCATTATGCTGTCCAAAGATCCCATACCTCCTATTGTGGGAAACAGTGGAAATCTAGCCAT
AGCTTACATGGATGTGTTTCGACCCAAAGTCCCAATCCATGTGGCTATGACGGGAGCCCTCAATGCTTGT
GGCGAGATTGAGAAAGTAAGCTTTAGAAGCACCAAGCTCGCTACTGCACACCGACTTGGCCTTAGGTTGG
CTGGTCCCGGAGCATTCGATGTAAACACCGGGCCCAACTGGGCAACGTTCATCAAACGTTTCCCTCACAA
TCCACGCGACTGGGACAGGCTCCCCTACCTCAACCTACCATACCTTCCACCCAATGCAGGACGCCAGTAC
CACCTTGCCATGGCTGCATCAGAGTTCAAAGAGACCCCCGAACTCGAGAGTGCCGTCAGAGCAATGGAAG
CAGCAGCCAACGTGGACCCACTATTCCAATCTGCACTCAGTGTGTTCATGTGGCTGGAAGAGAATGGGAT
TGTGACTGACATGGCCAACTTCGCACTCAGCGACCCGAACGCCCATCGGATGCGAAATTTTCTTGCAAAC
GCACCACAAGCAGGCAGCAAGTCGCAAAGGGCCAAGTACGGGACAGCAGGCTACGGAGTGGAGGCTCGGG
GCCCCACACCAGAGGAAGCACAGAGGGAAAAAGACACACGGATCTCAAAGAAGATGGAGACCATGGGCAT
CTACTTTGCAACACCAGAATGGGTAGCACTCAATGGGCACCGAGGGCCAAGCCCCGGCCAGCTAAAGTAC
TGGCAGAACACACGAGAAATACCGGACCCAAACGAGGACTATCTAGACTACGTGCATGCAGAGAAGAGCC
GGTTGGCATCAGAAGAACAAATCCTAAGGGCAGCTACGTCGATCTACGGGGCTCCAGGACAGGCAGAGCC
ACCCCAAGCTTTCATAGACGAAGTTGCCAAAGTCTATGAAATCAACCATGGACGTGGCCCAAACCAAGAA
CAGATGAAAGATCTGCTCTTGACTGCGATGGAGATGAAGCATCGCAATCCCAGGCGGGCTCTACCAAAGC
CCAAGCCAAAACCCAATGCTCCAACACAGAGACCTCCTGGTCGGCTGGGCCGCTGGATCAGGACCGTCTC
TGATGAGGACCTTGAGTGAGGCTCCTGGGAGTCTCCCGACACCACCCGCGCAGGTGTGGACACCAATTCG
GCCTTACAACATCCCAAATTGGATCCGTTCGCGGGTCCCCT

#### 1.2. IBDV WT Segment B

GGATACGATGGGTCTGACCCTCTGGGAGTCACGAATTAACGTGGCTACTAGGGGCGATACCCGCCGCTGG
CCGCCACGTTAGTGGCTCCTCTTCTTGATGATTCTGCCACCATGAGTGACGTTTTCAATAGTCCACAGGC
GCGAAGCACGATCTCAGCAGCGTTCGGCATAAAGCCTACTGCTGGACAAGACGTGGAAGAACTCTTGATC
CCTAAAGTTTGGGTGCCACCTGAGGATCCGCTTGCCAGCCCTAGTCGACTGGCAAAGTTCCTCAGAGAGA
ACGGCTACAAAGTTTTGCAGCCGCGGTCTCTGCCCGAGAATGAGGAGTATGAGACCGACCAAATACTCCC
AGACTTAGCATGGATGCGACAGATAGAAGGGGCTGTTTTAAAACCCACTCTATCTCTCCCTATTGGAGAT
CAGGAGTACTTCCCAAAGTACTACCCAACACATCGCCCTAGCAAGGAGAAGCCCAATGCGTACCCACCAG
ACATCGCACTACTCAAGCAGATGATTTACCTGTTTCTCCAGGTTCCAGAGGCCAACGAGGGCCTAAAGGA
TGAAGTAACCCTCTTGACCCAAAACATAAGGGACAAGGCCTATGGAAGTGGGACCTACATGGGACAAGCA
ACTCGACTTGTGGCCATGAAGGAGGTCGCCACTGGAAGAAACCCAAACAAGGATCCTCTAAAGCTTGGGT
ACACTTTTGAGAGCATCGCGCAGCTACTTGACATCACACTACCGGTAGGCCCACCCGGTGAGGATGACAA
GCCCTGGGTGCCACTCACAAGAGTGCCGTCACGGATGTTGGTGCTGACGGGAGACGTAGATGGCGACTTT
GAGGTTGAAGATTACCTTCCCAAAATCAACCTCAAGTCATCAAGTGGACTACCATATGTAGGTCGCACCA
AAGGAGAGACAATTGGCGAGATGATAGCTATATCAAACCAGTTTCTCAGAGAGCTATCAACACTGTTGAA
GCAAGGTGCAGGGACAAAGGGGTCAAACAAGAAGAAGCTACTCAGCATGTTAAGTGACTATTGGTACTTA
TCATGCGGGCTTTTGTTTCCAAAGGCTGAAAGGTACGACAAAAGTACATGGCTCACCAAGACCCGGAACA
TATGGTCAGCTCCATCCCCAACACACCTCATGATCTCCATGATCACCTGGCCCGTGATGTCCAACAGCCC
AAATAACGTGTTGAACATTGAAGGGTGTCCATCACTCTACAAATTCAACCCGTTCAGAGGAGGGTTGAAC
AGGATCGTCGAGTGGATATTGGCCCCGGAAGAACCCAAGGCTCTTGTATATGCGGACAACATATACATTG
TCCACTCAAACACGTGGTACTCAATTGACCTAGAGAAGGGTGAGGCAAACTGCACTCGCCAACACATGCA
AGCCGCAATGTACTACATACTCACCAGAGGGTGGTCAGACAACGGCGACCCAATGTTCAATCAAACATGG
GCCACCTTTGCCATGAACATTGCCCCTGCTCTAGTGGTGGACTCATCGTGCCTGATAATGAACCTGCAAA
TTAAGACCTATGGTCAAGGCAGCGGGAATGCAGCCACGTTCATCAACAACCACCTCTTGAGCACGCTAGT
GCTTGACCAGTGGAACTTGATGAGACAGCCCAGACCAGACAGCGAGGAGTTCAAATCAATTGAGGACAAG
CTAGGTATCAACTTTAAGATTGAGAGGTCCATTGATGATATCAGGGGCAAGCTGAGACAGCTTGTCCTCC
TTGCACAACCAGGGTACCTGAGTGGGGGGGTTGAACCAGAACAATCCAGCCCAACTGTTGAGCTTGACCT
ACTAGGGTGGTCAGCTACATACAGCAAAGATCTCGGGATCTATGTGCCGGTGCTTGACAAGGAACGCCTA
TTTTGTTCTGCTGCGTATCCCAAGGGAGTAGAGAACAAGAGTCTCAAGTCCAAAGTCGGGATCGAGCAGG
CATACAAGGTAGTCAGGTATGAGGCGTTGAGGTTGGTAGGTGGTTGGAACTACCCACTCCTGAACAAAGC
CTGCAAGAATAACGCAGGCGCCGCTCGGCGGCATCTGGAGGCCAAGGGGTTCCCACTCGACGAGTTCCTA
GCCGAGTGGTCTGAGCTGTCAGAGTTCGGTGAGGCCTTCGAAGGCTTCAATATCAAGCTGACCGTAACAT
CTGAGAGCCTAGCCGAACTGAACAAGCCAGTACCCCCCAAGCCCCCAAATGTCAACAGACCAGTCAACAC
TGGGGGACTCAAGGCAGTCAGCAACGCCCTCAAGACCGGTCGGTACAGGAACGAAGCCGGACTGAGTGGT
CTCGTCCTTCTAGCCACAGCAAGAAGCCGTCTGCAAGATGCAGTTAAGGCCAAGGCAGAAGCCGAGAAAC
TCCACAAGTCCAAGCCAGACGACCCCGATGCAGACTGGTTCGAAAGATCAGAAACTCTGTCAGACCTTCT
GGAGAAAGCCGACATCGCCAGCAAGGTCGCCCACTCAGCACTCGTGGAAACAAGCGACGCCCTTGAAGCA
GTTCAGTCGACTTCCGTGTACACCCCCAAGTACCCAGAAGTCAAGAACCCACAGACCGCCTCCAACCCCG
TTGTTGGGCTCCACCTGCCCGCCAAGAGAGCCACCGGTGTCCAGGCCGCTCTTCTCGGAGCAGGAACGAG
CAGACCAATGGGGATGGAGGCCCCAACACGGTCCAAGAACGCCGTGAAAATGGCCAAACGGCGGCAACGC
CAAAAGGAGAGCCGCTAACAGCCATGATGGGAACCACTCAAGAAGAGGACACTAATCCCAGACCCCGTAT
CCCCGGCCTTCGCCTGCGGGGGCCCCC

#### 1.3. IBDV P Segment A

GGATACGATCGGTCTGACCCCGAGGGAGTCACCCGGGGACAGGCCGTCAAGGCTTTGTTCCAGGATGGAA
CTCCTCCTTCTACAACGCTATCATTGATGGTTAGTAGAGATCAGACAAACGATCGCAGCGATGACAAACC
TGACAGATCAAACCCAACAGATTGTTCCGTTCATACGGAGCCTTCTGATGCCAACAACCGGACCGGCGTC
CATTCCGGACGACACCCTGGAGAAGCACACTCTCAGGTCAGAGACCTCGACCTACAATTTGACTGTGGGG
GACACAGGGTCAGGGCTAATTGTCTTTTTCCCTGGATTCCCTGGCTCAATTGTGGGTGCTCACTACACAC
TGCAGAGCAATGGGAACTACAAGTTCGATCAGATGCTCCTGACTGCCCAGAACCTACCGGCCAGTTACAA
CTACTGCAGGCTAGTGAGTCGGAGTCTCACAGTGAGGTCAAGCACACTCCCTGGTGGCGTTTATGCACTA
AACGGCACCATAAACGCCGTGACCTTCCAAGGAAGCCTGAGTGAACTGACAGATGTTAGCTACAATGGGT
TGATGTCTGCAACGGCCAACATCAACGACAAAATTGGGAATGTCCTGGTAGGGGAGGGGGTCACCGTCCT
CAGCTTACCCACATCATATGATCTTGGGTATGTGAGGCTTGGTGACCCCATTCCTGCTATAGGGCTTGAC

CCAAAAATGGTAGCCACATGTGACAGCAGTGACAGGCCCAGAGTCTACACCATAACTGCAGCCGATGATT
ACCAATTCTCATCACAGTACCAATCAGGTGGGGTAACAATCACACTGTTCTCAGCCAACATTGATGCTAT
CACAAGCCTCAGCATTGGGGGAGAGCTCGTGTTCCATACAAGCGTCCATGGCCTTGCACTGGACGCCACC
ATCTACCTTATAGGCTTTGATGGGACTACAGTAATCACCAGAGCTGTGGCCTCAGACAATGGGCTGACTA
CCGGCATCGACAATCTTATGCCATTCAATCTTGTGATTCCAACCAACGAGATAACCCAGCCAATCACATC
CATCAAACTGGAGATAGTGACCTCCAAAAGTGGCGGTCAGGCAGGGGACCAGATGTCATGGTCGGCAAGT
GGGAGCCTAGCAGTGACAATCCATGGTGGCAACTATCCAGGGGCCCTCCGTCCCGTCACACTAGTAGCCT
ACGAAAGAGTGGCAACAGGATCCGTCGTTACGGTCGCCGGGGTGAGCAACTTCGAGCTGATCCCAAATCC
TGAACTAGCAAAGAACCTGGTTACAGAATACGGCCGATTTGACCCAGGAGCCATGAACTACACAAAATTG
ATACTGAGTGAGAGGGACCGTCTTGGCATCAAGACCGTCTGGCCAACAAGGGAGTACACTGACTTTCGTG
AGTACTTCATGGAGGTGGCCGACCTCAACTCTCCCCTGAAGATTGCAGGAGCATTTGGCTTCAAAGACAT
AATCCGGGCCATAAGGAGGATAGCTGTGCCGGTGGTCTCTACATTGTTCCCACCTGCAGCTCCCCTAGCC
CTTGCAATTGGGGAAGGTGTAGACTACCTGCTGGGCGATGAGGCACAGGCTGCTTCAGGAACTGCTCGAG
CCGCGTCAGGAAAAGCAAGAGCTGCCTCAGGCCGTATAAGGCAGCTAACTCTCGCCGCTGACAAGGGGTA
CGAGGTTGTCGCGAATCTATTCCAGGTGCCCCAGAATCCCGTAGTCGACGGGATTCTTGCTTCACCTGGG
GTACTCCGCGGCGCACACAACCTCGACTGCGTGCTAAGAGAGGGTGCCACGCTATTCCCTGTGGTCATTA
CGACAGTGGAAGACGCCATGACACCCAAAGCACTGAACAGCAAAATGTTTGCTGTCATTGAAGGTGTGCG
AGAAGACCTCCAACCTCCATCTCAACGAGGATCCTTCATACGAACCCTCTCCGGACACAGAGTCTATGGA
TATGCTCCAGATGGGGTACTTCCACTGGAGACTGGGAGAGACTACACCGTTGTCCCAATAGATGATGTAT
GGGACGACAGCATTATGCTGTCCAAAGACCCCATACCTCCTATTGTGGGAAACAGTGGCAACCTAGCCAT
AGCTTACATGGATGTGTTTCGACCCAAAGTCCCCATCCATGTGGCCATGACAGGAGCCCTCAATGCTTGT
GGCGAGATTGAGAAAGTAAGCTTTAGAAGCACCAAGCTCGCCACTGCACACCGACTTGGCCTCAAGTTGG
CTGGTCCCGGAGCATTCGATGTAAACACCGGGCCCAATTGGGCAACGTTCATCAAACGTTTCCCTCACAA
TCCACGTGACTGGGACAGGCTCCCCTACCTCAACCTTCCATACCTTCCACCCAATGCAGGACGCCAGTAC
CACCTTGCCATGGCTGCATCAGAGTTCAAAGAGACCCCTGAACTCGAGAGCGCCGTCAGAGCCATGGAAG
CAGCAGCCAACGTGGACCCACTATTCCAATCTGCAATCAGTGTGTTCATGTGGCTGGAAGAGAATGGGAT
TGTGACTGACATGGCCAACTTCGCACTCAGCGACCCGAACGCCCATCGGATGCGAAATTTTCTTGCAAAC
GCACCACAAGCAGGCAGCAAGTCGCAACGGGCCAAGTACGGGACAGCAGGCTACGGAGTGGAGGCCCGGG
GCCCCACACCAGAGGAAGCGCAGAGGGAAAAAGACACACGGATCTCAAAGAAGATGGAGACCATGGGCAT
CTACTTTGCAACACCAGAATGGGTAGCACTCAATGGGCACCGAGGGCCAAGCCCGGGCCAGCTAAAGTAC
TGGCAGAACACACGAGAAATACCGGACCCAAACGAGGACTATCTAGACTACGTGCATGCAGAGAAGAGCC
GGTTGGCATCAGAAGAACAAATCCAAAGGGCAGCTACGTCGATCTACGGGGCTCCAGGACAGGCAGAGCC
ACCCCAAGCTTTCATAGACGAAGTTGCCAAAGTCTATGAAATCAACCATGGACGTGGCCCAAACCAAGAG
CAGATGAAAGATCTGCTCTTGACTGCGATGGAGATGAAGCATCGCAATCCCAGGCGGGCTCCACCAAAGC
CCAAGCCAAAACCCAATGCTCCAACACCGAGACCCCCTGGTCGGCTGGGCCGCTGGATCAGGACTGTCTC
TGATGAGGACCTTGAGTGAGGCTCCTGGGAGTCTCCCGACACCACCCGCGCAGGTGTGGACACCAATTCG
GTCTTACAACATCCCAAATTGGATCCGTTCGCGGGTCCCCT

#### 1.4. IBDV P Segment B

GGATACGATGGGTCTGACCCTCTGGGAGTCACGAATTAACATGGCTACTAGGGGCGATGCCCGCCTCTAA
CTGCCACGTTAGTGGCTCCTCTTCTTGATGATTCTGCCACCATGAGTGACATTTTCAACAGTCCACAGGC
GCGAAGCAAGATCTCAGCAGCGTTCGGCATAAAGCCTACTGCTGGACAAGACGTGGAAGAACTCTTGATC
CCTAAAGTCTGGGTGCCACCTGAGGATCCGCTTGCCAGCCCTAGTCGACTGGCAAAGTTCCTCAGAGAGA
ACGGCTACAAAGTTTTGCAGCCACGGTCTCTGCCCGAGAATGAGGAGTATGAGACCGACCAAATACTCCC
AGACTTAGCATGGATGCGACAGATAGAAGGGGCTGTTTTAAAACCTACTCTATCTCTCCCCATTGGAGAC
CAGGAGTACTTCCCAAAGTACTACCCAACACATCGCCCTAGCAAGGAGAAGACCAATGCGTACCCGCCAG
ACATCGCACTACTCAAGCAGATGATTTACCTGTTTCTCCAGGTTCCAGAGGCCAACGAGGGCCTAAAGGA
TGAAGTAACCCTCCTGACCCAAAATATAAGGGATAAGGCCTATGGAAGTGGGACCTACATGGGGCAAGCA
ACTCGACTTGTAGCCATGAAGGAGGTTGCCACTGGGAGAAACCCGAACAAGGATCCTCTAAAACTTGGGT
ACACTTTTGAGAGCATCGCGCAGCTGCTTGACATCACACTACCGGTAGGCCCACCCGGTGAGGATGACAA
GCCCTGGGTGCCACTCACAAGAGTGCCGTCGCGGATGTTGGTGCTGACGGGAGACGTAGATGGCGACTTT
GAGGTTGAAGATTACCTTCCCAAAATCAACCTCAAGTCATCAAGTGGACTACCATATGTAGGTCGCACCA
AAGGAGAGACAATTGGCGAGATGATAGCTATCTCAAACCAGTTTCTCAGAGAGCTATCAACACTGTTGAA
GCAAGGTGCAGGGACAAAGGGGTCAAACAAGAAGAAGCTACTCAGCATGTTAAGTGACTATTGGTACTTA
TCATGCGGGCTTTTGTTTCCAAAGGCTGAAAGGTACGACAAAAGCACATGGCTCACCAAAACCCGGAACA

```
TATGGTCAGCTCCATCCCCAACACACCTCATGATCTCCATGATCACCTGGCCCGTGATGTCCAACAGCCC
AAACAACGTGTTGAACATTGAAGGGTGTCCATCACTCTACAAATTCAACCCGTTCAGAGGAGGGTTGAAC
AGGATCGTCGAGTGGATATTGGCTCCGGAAGAACCCAAGGCTCTTGTATATGCGGACAACATATACATTG
TCCACTCAAACACGTGGTACTCAATTGACCTAGAGAAGGGCGAGGCAAACTGCACACGCCAACACATGCA
AGCCGCAATGTACTACATCCTCACCAGAGGGTGGTCAGACAACGGCGACCCAATGTTCAATCAAACATGG
GCCACCTTTGCAATGAACATTGCCCCTGCTCTAGTGGTAGACTCATCGTGCCTGATAATGAACCTGCAAA
TTAAGACCTATGGTCAAGGCAGCGGGAATGCAGCCACGTTCATCAATAACCACCTCTTGAGCACGCTAGT
GCTTGACCAGTGGAACCTGATGAGACAGCCCAGACCAGACAGCGAGGAGTTCAAATCAATTGAGGACAAG
CTAGGCATCAACTTCAAGATTGAGAGGTCCATTGATGACATCAGGGGCAAGCTGAGACAGCTTGTCCCCC
TTGCACAACCAGGGTACCTGAGTGGGGGGGTTGAACCAGAACAATCCAGCCCAACTGTTGAGCTTGACCT
ACTAGGGTGGTCAGCTACATACAGCAAAGATCTCGGGATCTATGTGCCGGTGCTTGACAAGGAACGCCTA
TTTTGTTCTGCGGCGTATCCCAAGGGAGTAGAGAACAAGAGTCTCAAGTCTAAAGTCGGGATCGAGCAGG
CATACAAGGTAGTCAGGTATGAGGCGTTGAGGTTGGTAGGTGGTTGGAACTACCCACTCCTGAACAAAGC
CTGCAAGAATAACGCAGGCGCTGCTCGGCGGCATCTGGAGGCCAAGGGGTTCCCACTCGACGAGTTCCTA
GCCGAGTGGTCCGAGCTGTCAGAGTTCGGTGAGGCCTTCGAAGGCTTCAATATCAAGCTGACCGTAACAT
CTGAGAGCCTAGCCGAACTGAACAAGCCAGTACCCCCCAAGCCCCCAAATGTCAACAGACCAGTCAACAC
TGGGGGACTCAAGGCAGTCAGCAACGCCCTTAAGACCGGTCGGTACAGGAACGAAGCCGGACTGAGTGGT
CTCGTCCTTCTAGCCACAGCAAGGAGCCGTCTGCAAGACGCAGTTAAGGCCAAGGCAGAAGCCGAGAAAC
TCCACAAGTCCAAGCCAGACGACCCCGATGCAGACTGGTTTGAAAGATCAGAAACTCTGTCAGACCTTCT
GGAGAAAGCCGACATCGCCAGTAAGGTCGCCCACTCAGCACTCGTGGAAACAAGCGACGCTCTTGAAGCA
GTTCAGTCGACTTCCGTGTACACCCCCAAGTACCCAGAAGTCAAGAACCCACAGACCGCCTCCAACCCCG
TTGTTGGGCTCCACCTGCCCGCCAAGAGAGCCACCGGTGTCCAGGCCGCTCTTCTCGGAGCAGGAACGAG
CCGACCAATGGGGATGGAGGCCCCAACACGGTCCAAGAACGCCGTGAAAATGGCCAAACGGCGGCAACGC
CAAAAAGAGAGCCGCCAATAGCCATGATGGGAACCACTCAAGAAGAGGACACTAATCCCAGACCCCGTAT
CCCCGGCCTTCGCCTGCGGGGGCCCCC
```

### 2. Genome Sequence Comparisons

Comparisons were performed with the LALIGN alignment tool (https://embnet.vital-it.ch/software/LALIGN_form.html) using default parameters.

#### 2.1. Segment A

```
              10        20        30        40        50        60
WT     GGAUACGAUCGGUCUGACCCGGGGGGAGUCACCCGGGGACAGGCUGACAAGGCCUUGUUC
       :::::::::::::::::::: : ::::::::::::::::::::: : :::::: ::::::
P      GGAUACGAUCGGUCUGACCCCGAGGGAGUCACCCGGGGACAGGCCGUCAAGGCUUUGUUC
              10        20        30        40        50        60

              70        80        90       100       110       120
WT     CAGGAUGGAACUCCUCCUUCUACAACGCUAUCAUUGAUGGUUAGUAGAGAUCAGACAAAC
       ::::::::::::::::::::::::::::::::::::::::::::::::::::::::::::
P      CAGGAUGGAACUCCUCCUUCUACAACGCUAUCAUUGAUGGUUAGUAGAGAUCAGACAAAC
              70        80        90       100       110       120

             130       140       150       160       170       180
WT     GAUCGCAGCGAUGACAAACCUGCAAGAUCAAACCCAACAGAUUGUUCCGUUCAUACGGAG
       ::::::::::::::::::::::  ::::::::::::::::::::::::::::::::::::
P      GAUCGCAGCGAUGACAAACCUGACAGAUCAAACCCAACAGAUUGUUCCGUUCAUACGGAG
             130       140       150       160       170       180

             190       200       210       220       230       240
WT     CCUUCUGAUGCCAACAACCGGACCGGCGUCCAUUCCGGACGACACCCUGGAGAAGCACAC
       ::::::::::::::::::::::::::::::::::::::::::::::::::::::::::::
P      CCUUCUGAUGCCAACAACCGGACCGGCGUCCAUUCCGGACGACACCCUGGAGAAGCACAC
             190       200       210       220       230       240

             250       260       270       280       290       300
WT     UCUCAGGUCAGAGACCUCGACCUACAAUUUGACUGUGGGGGACACAGGGUCAGGGCUAAU
       ::::::::::::::::::::::::::::::::::::::::::::::::::::::::::::
```

```
P       UCUCAGGUCAGAGACCUCGACCUACAAUUUGACUGUGGGGGACACAGGGUCAGGGCUAAU
               250       260       270       280       290       300

               310       320       330       340       350       360
WT      UGUCUUUUUCCCUGGAUUCCCUGGCUCAAUUGUGGGUGCUCACUACACACUGCAGGGCAA
        ::::::::::::::::::::::::::::::::::::::::::::::::::::::: ::::
P       UGUCUUUUUCCCUGGAUUCCCUGGCUCAAUUGUGGGUGCUCACUACACACUGCAGAGCAA
               310       320       330       340       350       360

               370       380       390       400       410       420
WT      UGGGAACUACAAGUUCGAUCAGAUGCUCCUGACUGCCCAGAACCUACCGGCCAGUUACAA
        ::::::::::::::::::::::::::::::::::::::::::::::::::::::::::::
P       UGGGAACUACAAGUUCGAUCAGAUGCUCCUGACUGCCCAGAACCUACCGGCCAGUUACAA
               370       380       390       400       410       420

               430       440       450       460       470       480
WT      CUACUGCAGGCUAGUGAGUCGGAGUCUCACAGUGAGGUCAAGCACACUUCCUGGUGGCGU
        :::::::::::::::::::::::::::::::::::::::::::::::: :::::::::::
P       CUACUGCAGGCUAGUGAGUCGGAGUCUCACAGUGAGGUCAAGCACACUCCCUGGUGGCGU
               430       440       450       460       470       480

               490       500       510       520       530       540
WT      UUAUGCACUAAACGGCACCAUAAACGCCGUGACCUUCCAAGGAAGCCUGAGUGAACUGAC
        ::::::::::::::::::::::::::::::::::::::::::::::::::::::::::::
P       UUAUGCACUAAACGGCACCAUAAACGCCGUGACCUUCCAAGGAAGCCUGAGUGAACUGAC
               490       500       510       520       530       540

               550       560       570       580       590       600
WT      AGAUGUUAGCUACAAUGGGUUGAUGUCUGCAACAGCCAACAUCAACGACAAAAUUGGGAA
        ::::::::::::::::::::::::::::::::: ::::::::::::::::::::::::::
P       AGAUGUUAGCUACAAUGGGUUGAUGUCUGCAACGGCCAACAUCAACGACAAAAUUGGGAA
               550       560       570       580       590       600

               610       620       630       640       650       660
WT      CGUCCUAGUAGGGGAAGGGGUCACCGUCCUCAGCUUACCCACAUCAUAUGAUCUUGGGUA
         ::::: :::::::: ::::::::::::::::::::::::::::::::::::::::::::
P       UGUCCUGGUAGGGGAGGGGGUCACCGUCCUCAGCUUACCCACAUCAUAUGAUCUUGGGUA
               610       620       630       640       650       660

               670       680       690       700       710       720
WT      UGUGAGGCUUGGUGACCCCAUUCCCGCAAUAGGGCUUGACCCAAAAAUGGUAGCCACAUG
        :::::::::::::::::::::::: :: ::::::::::::::::::::::::::::::::
P       UGUGAGGCUUGGUGACCCCAUUCCUGCUAUAGGGCUUGACCCAAAAAUGGUAGCCACAUG
               670       680       690       700       710       720

               730       740       750       760       770       780
WT      UGACAGCAGUGACAGGCCCAGAGUCUACACCAUAACUGCAGCCGAUGAUUACCAAUUCUC
        ::::::::::::::::::::::::::::::::::::::::::::::::::::::::::::
P       UGACAGCAGUGACAGGCCCAGAGUCUACACCAUAACUGCAGCCGAUGAUUACCAAUUCUC
               730       740       750       760       770       780

               790       800       810       820       830       840
WT      AUCACAGUACCAACCAGGUGGGGUAACAAUCACACUGUUCUCAGCCAACAUUGAUGCCAU
        ::::::::::::: ::::::::::::::::::::::::::::::::::::::::::: ::
P       AUCACAGUACCAAUCAGGUGGGGUAACAAUCACACUGUUCUCAGCCAACAUUGAUGCUAU
               790       800       810       820       830       840

               850       860       870       880       890       900
WT      CACAAGCCUCAGCGUUGGGGGAGAGCUCGUGUUUCGAACAAGCGUCCACGGCCUUGUACU
        ::::::::::::: ::::::::::::::::::: :  ::::::::::: ::::::: :::
P       CACAAGCCUCAGCAUUGGGGGAGAGCUCGUGUUCCAUACAAGCGUCCAUGGCCUUGCACU
               850       860       870       880       890       900

               910       920       930       940       950       960
WT      GGGCGCCACCAUCUACCUCAUAGGCUUUGAUGGGACAACGGUAAUCACCAGGGCUGUGGC
        :: ::::::::::::::: ::::::::::::::::: :: ::::::::::: ::::::::
P       GGACGCCACCAUCUACCUUAUAGGCUUUGAUGGGACUACAGUAAUCACCAGAGCUGUGGC
               910       920       930       940       950       960

               970       980       990      1000      1010      1020
WT      CGCAAACAAUGGGCUGACGACCGGCACCGACAACCUUAUGCCAUUCAAUCUUGUGAUUCC
        : :: ::::::::::::: ::::::: :::::: ::::::::::::::::::::::::::
P       CUCAGACAAUGGGCUGACUACCGGCAUCGACAAUCUUAUGCCAUUCAAUCUUGUGAUUCC
               970       980       990      1000      1010      1020

              1030      1040      1050      1060      1070      1080
WT      AACAAACGAGAUAACCCAGCCAAUCACAUCCAUCAAACUGGAGAUAGUGACCUCCAAAAG
```

```
          ::: ::::::::::::::::::::::::::::::::::::::::::::::::::::::::
P         AACCAACGAGAUAACCCAGCCAAUCACAUCCAUCAAACUGGAGAUAGUGACCUCCAAAAG
                1030      1040      1050      1060      1070      1080

                1090      1100      1110      1120      1130      1140
WT        UGGUGGUCAGGCAGGGGAUCAGAUGUCAUGGUCGGCAAGAGGGAGCCUAGCAGUGACGAU
          ::: :::::::::::::: :::::::::::::::::::: ::::::::::::::::: ::
P         UGGCGGUCAGGCAGGGGACCAGAUGUCAUGGUCGGCAAGUGGGAGCCUAGCAGUGACAAU
                1090      1100      1110      1120      1130      1140

                1150      1160      1170      1180      1190      1200
WT        CCAUGGUGGCAACUAUCCAGGGGCCCUCCGUCCCGUCACGCUAGUGGCCUACGAAAGAGU
          ::::::::::::::::::::::::::::::::::::::: ::::: ::::::::::::::
P         CCAUGGUGGCAACUAUCCAGGGGCCCUCCGUCCCGUCACACUAGUAGCCUACGAAAGAGU
                1150      1160      1170      1180      1190      1200

                1210      1220      1230      1240      1250      1260
WT        GGCAACAGGAUCCGUCGUUACGGUCGCUGGGGUGAGCAACUUCGAGCUGAUCCCAAAUCC
          ::::::::::::::::::::::::::: ::::::::::::::::::::::::::::::::
P         GGCAACAGGAUCCGUCGUUACGGUCGCCGGGGUGAGCAACUUCGAGCUGAUCCCAAAUCC
                1210      1220      1230      1240      1250      1260

                1270      1280      1290      1300      1310      1320
WT        UGAACUAGCAAAGAACCUGGUUACAGAAUACGGCCGAUUUGACCCAGGAGCCAUGAACUA
          ::::::::::::::::::::::::::::::::::::::::::::::::::::::::::::
P         UGAACUAGCAAAGAACCUGGUUACAGAAUACGGCCGAUUUGACCCAGGAGCCAUGAACUA
                1270      1280      1290      1300      1310      1320

                1330      1340      1350      1360      1370      1380
WT        CACAAAAUUGAUACUGAGUGAGAGGGACCGUCUUGGCAUCAAGACCGUCUGGCCAACAAG
          ::::::::::::::::::::::::::::::::::::::::::::::::::::::::::::
P         CACAAAAUUGAUACUGAGUGAGAGGGACCGUCUUGGCAUCAAGACCGUCUGGCCAACAAG
                1330      1340      1350      1360      1370      1380

                1390      1400      1410      1420      1430      1440
WT        GGAGUACACUGACUUUCGUGAAUACUUCAUGGAGGUGGCCGACCUCAACUCUCCCCUGAA
          ::::::::::::::::::::: ::::::::::::::::::::::::::::::::::::::
P         GGAGUACACUGACUUUCGUGAGUACUUCAUGGAGGUGGCCGACCUCAACUCUCCCCUGAA
                1390      1400      1410      1420      1430      1440

                1450      1460      1470      1480      1490      1500
WT        GAUUGCAGGAGCAUUCGGCUUCAAAGACAUAAUCCGGGCCAUAAGGAGGAUAGCUGUGCC
          ::::::::::::::: ::::::::::::::::::::::::::::::::::::::::::::
P         GAUUGCAGGAGCAUUUGGCUUCAAAGACAUAAUCCGGGCCAUAAGGAGGAUAGCUGUGCC
                1450      1460      1470      1480      1490      1500

                1510      1520      1530      1540      1550      1560
WT        GGUGGUCUCCACAUUGUUCCCACCUGCCGCUCCCCUAGCCCAUGCAAUUGGGGAAGGUGU
          ::::::::: ::::::::::::::::: ::::::::::::: ::::::::::::::::::
P         GGUGGUCUCUACAUUGUUCCCACCUGCAGCUCCCCUAGCCCUUGCAAUUGGGGAAGGUGU
                1510      1520      1530      1540      1550      1560

                1570      1580      1590      1600      1610      1620
WT        AGACUACCUGCUGGGCGAUGAGGACCAGGCCGCUUCAGGAACUGCUCGAGCCGCGUCAGG
          :::::::::::::::::::::::  ::::: :::::::::::::::::::::::::::::
P         AGACUACCUGCUGGGCGAUGAGGCACAGGCUGCUUCAGGAACUGCUCGAGCCGCGUCAGG
                1570      1580      1590      1600      1610      1620

                1630      1640      1650      1660      1670      1680
WT        AAAAGCAAGAGCUGCCUCAGGCCGCAUAAGGCAGCUGACUCUCGCCGCCGACAAGGGGUA
          :::::::::::::::::::::::: ::::::::::: ::::::::::: :::::::::::
P         AAAAGCAAGAGCUGCCUCAGGCCGUAUAAGGCAGCUAACUCUCGCCGCUGACAAGGGGUA
                1630      1640      1650      1660      1670      1680

                1690      1700      1710      1720      1730      1740
WT        CGAGGUAGUCGCGAAUCUAUUCCAGGUGCCCCAGAAUCCCGUAGUCGACGGGAUUCUUGC
          :::::: :::::::::::::::::::::::::::::::::::::::::::::::::::::
P         CGAGGUUGUCGCGAAUCUAUUCCAGGUGCCCCAGAAUCCCGUAGUCGACGGGAUUCUUGC
                1690      1700      1710      1720      1730      1740

                1750      1760      1770      1780      1790      1800
WT        UUCACCUGGGGUACUCCGCGGUGCACACAACCUCGACUGCGUGUUAAGAGAGGGUGCCAC
          ::::::::::::::::::::: ::::::::::::::::::::: ::::::::::::::::
P         UUCACCUGGGGUACUCCGCGGCGCACACAACCUCGACUGCGUGCUAAGAGAGGGUGCCAC
                1750      1760      1770      1780      1790      1800

                1810      1820      1830      1840      1850      1860
```

```
WT     GCUAUUCCCUGUGGUUAUUACGACAGUGGAAGACGCCAUGACACCCAAAGCAUUGAACAG
       ::::::::::::::: :::::::::::::::::::::::::::::::::::: :::::::
P      GCUAUUCCCUGUGGUCAUUACGACAGUGGAAGACGCCAUGACACCCAAAGCACUGAACAG
             1810      1820      1830      1840      1850      1860

             1870      1880      1890      1900      1910      1920
WT     CAAAAUGUUUGCUGUCAUUGAAGGCGUGCGAGAAGACCUCCAACCUCCAUCUCAAAGAGG
       :::::::::::::::::::::::: :::::::::::::::::::::::::::::: ::::
P      CAAAAUGUUUGCUGUCAUUGAAGGUGUGCGAGAAGACCUCCAACCUCCAUCUCAACGAGG
             1870      1880      1890      1900      1910      1920

             1930      1940      1950      1960      1970      1980
WT     AUCCUUCAUACGAACUCUCUCUGGACACAGAGUCUAUGGAUAUGCUCCAGAUGGGGUACU
       ::::::::::::::: ::::: ::::::::::::::::::::::::::::::::::::::
P      AUCCUUCAUACGAACCCUCUCCGGACACAGAGUCUAUGGAUAUGCUCCAGAUGGGGUACU
             1930      1940      1950      1960      1970      1980

             1990      2000      2010      2020      2030      2040
WT     UCCACUGGAGACUGGGAGAGACUACACCGUUGUCCCAAUAGAUGAUGUCUGGGACGACAG
       :::::::::::::::::::::::::::::::::::::::::::::::: :::::::::::
P      UCCACUGGAGACUGGGAGAGACUACACCGUUGUCCCAAUAGAUGAUGUAUGGGACGACAG
             1990      2000      2010      2020      2030      2040

             2050      2060      2070      2080      2090      2100
WT     CAUUAUGCUGUCCAAAGAUCCCAUACCUCCUAUUGUGGGAAACAGUGGAAAUCUAGCCAU
       :::::::::::::::::: ::::::::::::::::::::::::::::: :: ::::::::
P      CAUUAUGCUGUCCAAAGACCCCAUACCUCCUAUUGUGGGAAACAGUGGCAACCUAGCCAU
             2050      2060      2070      2080      2090      2100

             2110      2120      2130      2140      2150      2160
WT     AGCUUACAUGGAUGUGUUUCGACCCAAAGUCCCAAUCCAUGUGGCUAUGACGGGAGCCCU
       ::::::::::::::::::::::::::::::::: ::::::::::: ::::: ::::::::
P      AGCUUACAUGGAUGUGUUUCGACCCAAAGUCCCCAUCCAUGUGGCCAUGACAGGAGCCCU
             2110      2120      2130      2140      2150      2160

             2170      2180      2190      2200      2210      2220
WT     CAAUGCUUGUGGCGAGAUUGAGAAAGUAAGCUUUAGAAGCACCAAGCUCGCUACUGCACA
       ::::::::::::::::::::::::::::::::::::::::::::::::::: ::::::::
P      CAAUGCUUGUGGCGAGAUUGAGAAAGUAAGCUUUAGAAGCACCAAGCUCGCCACUGCACA
             2170      2180      2190      2200      2210      2220

             2230      2240      2250      2260      2270      2280
WT     CCGACUUGGCCUUAGGUUGGCUGGUCCCGGAGCAUUCGAUGUAAACACCGGGCCCAACUG
       :::::::::::: : :::::::::::::::::::::::::::::::::::::::::: ::
P      CCGACUUGGCCUCAAGUUGGCUGGUCCCGGAGCAUUCGAUGUAAACACCGGGCCCAAUUG
             2230      2240      2250      2260      2270      2280

             2290      2300      2310      2320      2330      2340
WT     GGCAACGUUCAUCAAACGUUUCCCUCACAAUCCACGCGACUGGGACAGGCUCCCCUACCU
       :::::::::::::::::::::::::::::::::::: :::::::::::::::::::::::
P      GGCAACGUUCAUCAAACGUUUCCCUCACAAUCCACGUGACUGGGACAGGCUCCCCUACCU
             2290      2300      2310      2320      2330      2340

             2350      2360      2370      2380      2390      2400
WT     CAACCUACCAUACCUUCCACCCAAUGCAGGACGCCAGUACCACCUUGCCAUGGCUGCAUC
       :::::: :::::::::::::::::::::::::::::::::::::::::::::::::::::
P      CAACCUUCCAUACCUUCCACCCAAUGCAGGACGCCAGUACCACCUUGCCAUGGCUGCAUC
             2350      2360      2370      2380      2390      2400

             2410      2420      2430      2440      2450      2460
WT     AGAGUUCAAAGAGACCCCCGAACUCGAGAGUGCCGUCAGAGCAAUGGAAGCAGCAGCCAA
       :::::::::::::::::: ::::::::::: ::::::::::: :::::::::::::::::
P      AGAGUUCAAAGAGACCCCUGAACUCGAGAGCGCCGUCAGAGCCAUGGAAGCAGCAGCCAA
             2410      2420      2430      2440      2450      2460

             2470      2480      2490      2500      2510      2520
WT     CGUGGACCCACUAUUCCAAUCUGCACUCAGUGUGUUCAUGUGGCUGGAAGAGAAUGGGAU
       ::::::::::::::::::::::::: ::::::::::::::::::::::::::::::::::
P      CGUGGACCCACUAUUCCAAUCUGCAAUCAGUGUGUUCAUGUGGCUGGAAGAGAAUGGGAU
             2470      2480      2490      2500      2510      2520

             2530      2540      2550      2560      2570      2580
WT     UGUGACUGACAUGGCCAACUUCGCACUCAGCGACCCGAACGCCCAUCGGAUGCGAAAUUU
       ::::::::::::::::::::::::::::::::::::::::::::::::::::::::::::
P      UGUGACUGACAUGGCCAACUUCGCACUCAGCGACCCGAACGCCCAUCGGAUGCGAAAUUU
             2530      2540      2550      2560      2570      2580
```

```
          2590      2600      2610      2620      2630      2640
WT   UCUUGCAAACGCACCACAAGCAGGCAGCAAGUCGCAAAGGGCCAAGUACGGGACAGCAGG
     ::::::::::::::::::::::::::::::::::::: ::::::::::::::::::::::
P    UCUUGCAAACGCACCACAAGCAGGCAGCAAGUCGCAACGGGCCAAGUACGGGACAGCAGG
          2590      2600      2610      2620      2630      2640

          2650      2660      2670      2680      2690      2700
WT   CUACGGAGUGGAGGCUCGGGGCCCCACACCAGAGGAAGCACAGAGGGAAAAAGACACACG
     ::::::::::::::: ::::::::::::::::::::::: ::::::::::::::::::::
P    CUACGGAGUGGAGGCCCGGGGCCCCACACCAGAGGAAGCGCAGAGGGAAAAAGACACACG
          2650      2660      2670      2680      2690      2700

          2710      2720      2730      2740      2750      2760
WT   GAUCUCAAAGAAGAUGGAGACCAUGGGCAUCUACUUUGCAACACCAGAAUGGGUAGCACU
     ::::::::::::::::::::::::::::::::::::::::::::::::::::::::::::
P    GAUCUCAAAGAAGAUGGAGACCAUGGGCAUCUACUUUGCAACACCAGAAUGGGUAGCACU
          2710      2720      2730      2740      2750      2760

          2770      2780      2790      2800      2810      2820
WT   CAAUGGGCACCGAGGGCCAAGCCCCGGCCAGCUAAAGUACUGGCAGAACACACGAGAAAU
     :::::::::::::::::::::::: :::::::::::::::::::::::::::::::::::
P    CAAUGGGCACCGAGGGCCAAGCCCGGGCCAGCUAAAGUACUGGCAGAACACACGAGAAAU
          2770      2780      2790      2800      2810      2820

          2830      2840      2850      2860      2870      2880
WT   ACCGGACCCAAACGAGGACUAUCUAGACUACGUGCAUGCAGAGAAGAGCCGGUUGGCAUC
     ::::::::::::::::::::::::::::::::::::::::::::::::::::::::::::
P    ACCGGACCCAAACGAGGACUAUCUAGACUACGUGCAUGCAGAGAAGAGCCGGUUGGCAUC
          2830      2840      2850      2860      2870      2880

          2890      2900      2910      2920      2930      2940
WT   AGAAGAACAAAUCCUAAGGGCAGCUACGUCGAUCUACGGGGCUCCAGGACAGGCAGAGCC
     :::::::::::::: :::::::::::::::::::::::::::::::::::::::::::::
P    AGAAGAACAAAUCCAAAGGGCAGCUACGUCGAUCUACGGGGCUCCAGGACAGGCAGAGCC
          2890      2900      2910      2920      2930      2940

          2950      2960      2970      2980      2990      3000
WT   ACCCCAAGCUUUCAUAGACGAAGUUGCCAAAGUCUAUGAAAUCAACCAUGGACGUGGCCC
     ::::::::::::::::::::::::::::::::::::::::::::::::::::::::::::
P    ACCCCAAGCUUUCAUAGACGAAGUUGCCAAAGUCUAUGAAAUCAACCAUGGACGUGGCCC
          2950      2960      2970      2980      2990      3000

          3010      3020      3030      3040      3050      3060
WT   AAACCAAGAACAGAUGAAAGAUCUGCUCUUGACUGCGAUGGAGAUGAAGCAUCGCAAUCC
     ::::::::: ::::::::::::::::::::::::::::::::::::::::::::::::::
P    AAACCAAGAGCAGAUGAAAGAUCUGCUCUUGACUGCGAUGGAGAUGAAGCAUCGCAAUCC
          3010      3020      3030      3040      3050      3060

          3070      3080      3090      3100      3110      3120
WT   CAGGCGGGCUCUACCAAAGCCCAAGCCAAAACCCAAUGCUCCAACACAGAGACCUCCUGG
     ::::::::::: ::::::::::::::::::::::::::::::::::: :::::: :::::
P    CAGGCGGGCUCCACCAAAGCCCAAGCCAAAACCCAAUGCUCCAACACCGAGACCCCCUGG
          3070      3080      3090      3100      3110      3120

          3130      3140      3150      3160      3170      3180
WT   UCGGCUGGGCCGCUGGAUCAGGACCGUCUCUGAUGAGGACCUUGAGUGAGGCUCCUGGGA
     :::::::::::::::::::::::: :::::::::::::::::::::::::::::::::::
P    UCGGCUGGGCCGCUGGAUCAGGACUGUCUCUGAUGAGGACCUUGAGUGAGGCUCCUGGGA
          3130      3140      3150      3160      3170      3180

          3190      3200      3210      3220      3230      3240
WT   GUCUCCCGACACCACCCGCGCAGGUGUGGACACCAAUUCGGCCUUACAACAUCCCAAAUU
     ::::::::::::::::::::::::::::::::::::::::: ::::::::::::::::::
P    GUCUCCCGACACCACCCGCGCAGGUGUGGACACCAAUUCGGUCUUACAACAUCCCAAAUU
          3190      3200      3210      3220      3230      3240

          3250      3260
WT   GGAUCCGUUCGCGGGUCCCCU
     :::::::::::::::::::::
P    GGAUCCGUUCGCGGGUCCCCU
          3250      3260
```

97.3% identity in 3,261 nt overlap.

### 2.2. Segment B

```
                10        20        30        40        50        60
WT      GGAUACGAUGGGUCUGACCCUCUGGGAGUCACGAAUUAACGUGGCUACUAGGGGCGAUAC
        :::::::::::::::::::::::::::::::::::::::: ::::::::::::::::: :
P       GGAUACGAUGGGUCUGACCCUCUGGGAGUCACGAAUUAACAUGGCUACUAGGGGCGAUGC
                10        20        30        40        50        60

                70        80        90       100       110       120
WT      CCGCCGCUGGCCGCCACGUUAGUGGCUCCUCUUCUUGAUGAUUCUGCCACCAUGAGUGAC
        ::::: ::  : ::::::::::::::::::::::::::::::::::::::::::::::::
P       CCGCCUCUAACUGCCACGUUAGUGGCUCCUCUUCUUGAUGAUUCUGCCACCAUGAGUGAC
                70        80        90       100       110       120

               130       140       150       160       170       180
WT      GUUUUCAAUAGUCCACAGGCGCGAAGCACGAUCUCAGCAGCGUUCGGCAUAAAGCCUACU
         ::::::: ::::::::::::::::::: :::::::::::::::::::::::::::::::
P       AUUUUCAACAGUCCACAGGCGCGAAGCAAGAUCUCAGCAGCGUUCGGCAUAAAGCCUACU
               130       140       150       160       170       180

               190       200       210       220       230       240
WT      GCUGGACAAGACGUGGAAGAACUCUUGAUCCCUAAAGUUUGGGUGCCACCUGAGGAUCCG
        :::::::::::::::::::::::::::::::::::::: :::::::::::::::::::::
P       GCUGGACAAGACGUGGAAGAACUCUUGAUCCCUAAAGUCUGGGUGCCACCUGAGGAUCCG
               190       200       210       220       230       240

               250       260       270       280       290       300
WT      CUUGCCAGCCCUAGUCGACUGGCAAAGUUCCUCAGAGAGAACGGCUACAAAGUUUUGCAG
        ::::::::::::::::::::::::::::::::::::::::::::::::::::::::::::
P       CUUGCCAGCCCUAGUCGACUGGCAAAGUUCCUCAGAGAGAACGGCUACAAAGUUUUGCAG
               250       260       270       280       290       300

               310       320       330       340       350       360
WT      CCGCGGUCUCUGCCCGAGAAUGAGGAGUAUGAGACCGACCAAAUACUCCCAGACUUAGCA
        :: :::::::::::::::::::::::::::::::::::::::::::::::::::::::::
P       CCACGGUCUCUGCCCGAGAAUGAGGAGUAUGAGACCGACCAAAUACUCCCAGACUUAGCA
               310       320       330       340       350       360

               370       380       390       400       410       420
WT      UGGAUGCGACAGAUAGAAGGGGCUGUUUUAAAACCCACUCUAUCUCUCCCUAUUGGAGAU
        ::::::::::::::::::::::::::::::::::: :::::::::::::: :::::::: 
P       UGGAUGCGACAGAUAGAAGGGGCUGUUUUAAAACCUACUCUAUCUCUCCCCAUUGGAGAC
               370       380       390       400       410       420

               430       440       450       460       470       480
WT      CAGGAGUACUUCCCAAAGUACUACCCAACACAUCGCCCUAGCAAGGAGAAGCCCAAUGCG
        ::::::::::::::::::::::::::::::::::::::::::::::::::: ::::::::
P       CAGGAGUACUUCCCAAAGUACUACCCAACACAUCGCCCUAGCAAGGAGAAGACCAAUGCG
               430       440       450       460       470       480

               490       500       510       520       530       540
WT      UACCCACCAGACAUCGCACUACUCAAGCAGAUGAUUUACCUGUUUCUCCAGGUUCCAGAG
        ::::: ::::::::::::::::::::::::::::::::::::::::::::::::::::::
P       UACCCGCCAGACAUCGCACUACUCAAGCAGAUGAUUUACCUGUUUCUCCAGGUUCCAGAG
               490       500       510       520       530       540

               550       560       570       580       590       600
WT      GCCAACGAGGGCCUAAAGGAUGAAGUAACCCUCUUGACCCAAAACAUAAGGGACAAGGCC
        ::::::::::::::::::::::::::::::::: :::::::::: :::::::: ::::::
P       GCCAACGAGGGCCUAAAGGAUGAAGUAACCCUCCUGACCCAAAAUAUAAGGGAUAAGGCC
               550       560       570       580       590       600

               610       620       630       640       650       660
WT      UAUGGAAGUGGGACCUACAUGGGACAAGCAACUCGACUUGUGGCCAUGAAGGAGGUCGCC
        ::::::::::::::::::::::: ::::::::::::::::: :::::::::::::: :::
P       UAUGGAAGUGGGACCUACAUGGGGCAAGCAACUCGACUUGUAGCCAUGAAGGAGGUUGCC
               610       620       630       640       650       660

               670       680       690       700       710       720
WT      ACUGGAAGAAACCCAAACAAGGAUCCUCUAAAGCUUGGGUACACUUUUGAGAGCAUCGCG
        ::::: :::::::: ::::::::::::::::: :::::::::::::::::::::::::::
```

```
P      ACUGGGAGAAACCCGAACAAGGAUCCUCUAAAACUUGGGUACACUUUUGAGAGCAUCGCG
              670       680       690       700       710       720

              730       740       750       760       770       780
WT     CAGCUACUUGACAUCACACUACCGGUAGGCCCACCCGGUGAGGAUGACAAGCCCUGGGUG
       ::::: ::::::::::::::::::::::::::::::::::::::::::::::::::::::
P      CAGCUGCUUGACAUCACACUACCGGUAGGCCCACCCGGUGAGGAUGACAAGCCCUGGGUG
              730       740       750       760       770       780

              790       800       810       820       830       840
WT     CCACUCACAAGAGUGCCGUCACGGAUGUUGGUGCUGACGGGAGACGUAGAUGGCGACUUU
       :::::::::::::::::::: :::::::::::::::::::::::::::::::::::::::
P      CCACUCACAAGAGUGCCGUCGCGGAUGUUGGUGCUGACGGGAGACGUAGAUGGCGACUUU
              790       800       810       820       830       840

              850       860       870       880       890       900
WT     GAGGUUGAAGAUUACCUUCCCAAAAUCAACCUCAAGUCAUCAAGUGGACUACCAUAUGUA
       ::::::::::::::::::::::::::::::::::::::::::::::::::::::::::::
P      GAGGUUGAAGAUUACCUUCCCAAAAUCAACCUCAAGUCAUCAAGUGGACUACCAUAUGUA
              850       860       870       880       890       900

              910       920       930       940       950       960
WT     GGUCGCACCAAAGGAGAGACAAUUGGCGAGAUGAUAGCUAUAUCAAACCAGUUUCUCAGA
       ::::::::::::::::::::::::::::::::::::::::: ::::::::::::::::::
P      GGUCGCACCAAAGGAGAGACAAUUGGCGAGAUGAUAGCUAUCUCAAACCAGUUUCUCAGA
              910       920       930       940       950       960

              970       980       990      1000      1010      1020
WT     GAGCUAUCAACACUGUUGAAGCAAGGUGCAGGGACAAAGGGGUCAAACAAGAAGAAGCUA
       ::::::::::::::::::::::::::::::::::::::::::::::::::::::::::::
P      GAGCUAUCAACACUGUUGAAGCAAGGUGCAGGGACAAAGGGGUCAAACAAGAAGAAGCUA
              970       980       990      1000      1010      1020

             1030      1040      1050      1060      1070      1080
WT     CUCAGCAUGUUAAGUGACUAUUGGUACUUAUCAUGCGGGCUUUUGUUUCCAAAGGCUGAA
       ::::::::::::::::::::::::::::::::::::::::::::::::::::::::::::
P      CUCAGCAUGUUAAGUGACUAUUGGUACUUAUCAUGCGGGCUUUUGUUUCCAAAGGCUGAA
             1030      1040      1050      1060      1070      1080

             1090      1100      1110      1120      1130      1140
WT     AGGUACGACAAAAGUACAUGGCUCACCAAGACCCGGAACAUAUGGUCAGCUCCAUCCCCA
       :::::::::::::: :::::::::::::: ::::::::::::::::::::::::::::::
P      AGGUACGACAAAAGCACAUGGCUCACCAAAACCCGGAACAUAUGGUCAGCUCCAUCCCCA
             1090      1100      1110      1120      1130      1140

             1150      1160      1170      1180      1190      1200
WT     ACACACCUCAUGAUCUCCAUGAUCACCUGGCCCGUGAUGUCCAACAGCCCAAAUAACGUG
       ::::::::::::::::::::::::::::::::::::::::::::::::::::: ::::::
P      ACACACCUCAUGAUCUCCAUGAUCACCUGGCCCGUGAUGUCCAACAGCCCAAACAACGUG
             1150      1160      1170      1180      1190      1200

             1210      1220      1230      1240      1250      1260
WT     UUGAACAUUGAAGGGUGUCCAUCACUCUACAAAUUCAACCCGUUCAGAGGAGGGUUGAAC
       ::::::::::::::::::::::::::::::::::::::::::::::::::::::::::::
P      UUGAACAUUGAAGGGUGUCCAUCACUCUACAAAUUCAACCCGUUCAGAGGAGGGUUGAAC
             1210      1220      1230      1240      1250      1260

             1270      1280      1290      1300      1310      1320
WT     AGGAUCGUCGAGUGGAUAUUGGCCCCGGAAGAACCCAAGGCUCUUGUAUAUGCGGACAAC
       ::::::::::::::::::::::: ::::::::::::::::::::::::::::::::::::
P      AGGAUCGUCGAGUGGAUAUUGGCUCCGGAAGAACCCAAGGCUCUUGUAUAUGCGGACAAC
             1270      1280      1290      1300      1310      1320

             1330      1340      1350      1360      1370      1380
WT     AUAUACAUUGUCCACUCAAACACGUGGUACUCAAUUGACCUAGAGAAGGGUGAGGCAAAC
       :::::::::::::::::::::::::::::::::::::::::::::::::: :::::::::
P      AUAUACAUUGUCCACUCAAACACGUGGUACUCAAUUGACCUAGAGAAGGGCGAGGCAAAC
             1330      1340      1350      1360      1370      1380

             1390      1400      1410      1420      1430      1440
WT     UGCACUCGCCAACACAUGCAAGCCGCAAUGUACUACAUACUCACCAGAGGGUGGUCAGAC
       ::::: :::::::::::::::::::::::::::::::: :::::::::::::::::::::
P      UGCACACGCCAACACAUGCAAGCCGCAAUGUACUACAUCCUCACCAGAGGGUGGUCAGAC
             1390      1400      1410      1420      1430      1440

             1450      1460      1470      1480      1490      1500
WT     AACGGCGACCCAAUGUUCAAUCAAACAUGGGCCACCUUUGCCAUGAACAUUGCCCCUGCU
```

```
          ::::::::::::::::::::::::::::::::::::::::: ::::::::::::::::::
P         AACGGCGACCCAAUGUUCAAUCAAACAUGGGCCACCUUUGCAAUGAACAUUGCCCCUGCU
                1450      1460      1470      1480      1490      1500

                1510      1520      1530      1540      1550      1560
WT        CUAGUGGUGGACUCAUCGUGCCUGAUAAUGAACCUGCAAAUUAAGACCUAUGGUCAAGGC
          :::::::: :::::::::::::::::::::::::::::::::::::::::::::::::::
P         CUAGUGGUAGACUCAUCGUGCCUGAUAAUGAACCUGCAAAUUAAGACCUAUGGUCAAGGC
                1510      1520      1530      1540      1550      1560

                1570      1580      1590      1600      1610      1620
WT        AGCGGGAAUGCAGCCACGUUCAUCAACAACCACCUCUUGAGCACGCUAGUGCUUGACCAG
          ::::::::::::::::::::::::::: ::::::::::::::::::::::::::::::::
P         AGCGGGAAUGCAGCCACGUUCAUCAAUAACCACCUCUUGAGCACGCUAGUGCUUGACCAG
                1570      1580      1590      1600      1610      1620

                1630      1640      1650      1660      1670      1680
WT        UGGAACUUGAUGAGACAGCCCAGACCAGACAGCGAGGAGUUCAAAUCAAUUGAGGACAAG
          :::::: :::::::::::::::::::::::::::::::::::::::::::::::::::::
P         UGGAACCUGAUGAGACAGCCCAGACCAGACAGCGAGGAGUUCAAAUCAAUUGAGGACAAG
                1630      1640      1650      1660      1670      1680

                1690      1700      1710      1720      1730      1740
WT        CUAGGUAUCAACUUUAAGAUUGAGAGGUCCAUUGAUGAUAUCAGGGGCAAGCUGAGACAG
          ::::: :::::::: ::::::::::::::::::::::: :::::::::::::::::::::
P         CUAGGCAUCAACUUCAAGAUUGAGAGGUCCAUUGAUGACAUCAGGGGCAAGCUGAGACAG
                1690      1700      1710      1720      1730      1740

                1750      1760      1770      1780      1790      1800
WT        CUUGUCCUCCUUGCACAACCAGGGUACCUGAGUGGGGGGGUUGAACCAGAACAAUCCAGC
          ::::::: ::::::::::::::::::::::::::::::::::::::::::::::::::::
P         CUUGUCCCCCUUGCACAACCAGGGUACCUGAGUGGGGGGGUUGAACCAGAACAAUCCAGC
                1750      1760      1770      1780      1790      1800

                1810      1820      1830      1840      1850      1860
WT        CCAACUGUUGAGCUUGACCUACUAGGGUGGUCAGCUACAUACAGCAAAGAUCUCGGGAUC
          ::::::::::::::::::::::::::::::::::::::::::::::::::::::::::::
P         CCAACUGUUGAGCUUGACCUACUAGGGUGGUCAGCUACAUACAGCAAAGAUCUCGGGAUC
                1810      1820      1830      1840      1850      1860

                1870      1880      1890      1900      1910      1920
WT        UAUGUGCCGGUGCUUGACAAGGAACGCCUAUUUUGUUCUGCUGCGUAUCCCAAGGGAGUA
          ::::::::::::::::::::::::::::::::::::::::: ::::::::::::::::::
P         UAUGUGCCGGUGCUUGACAAGGAACGCCUAUUUUGUUCUGCGGCGUAUCCCAAGGGAGUA
                1870      1880      1890      1900      1910      1920

                1930      1940      1950      1960      1970      1980
WT        GAGAACAAGAGUCUCAAGUCCAAAGUCGGGAUCGAGCAGGCAUACAAGGUAGUCAGGUAU
          :::::::::::::::::::: :::::::::::::::::::::::::::::::::::::::
P         GAGAACAAGAGUCUCAAGUCUAAAGUCGGGAUCGAGCAGGCAUACAAGGUAGUCAGGUAU
                1930      1940      1950      1960      1970      1980

                1990      2000      2010      2020      2030      2040
WT        GAGGCGUUGAGGUUGGUAGGUGGUUGGAACUACCCACUCCUGAACAAAGCCUGCAAGAAU
          ::::::::::::::::::::::::::::::::::::::::::::::::::::::::::::
P         GAGGCGUUGAGGUUGGUAGGUGGUUGGAACUACCCACUCCUGAACAAAGCCUGCAAGAAU
                1990      2000      2010      2020      2030      2040

                2050      2060      2070      2080      2090      2100
WT        AACGCAGGCGCCGCUCGGCGGCAUCUGGAGGCCAAGGGGUUCCCACUCGACGAGUUCCUA
          ::::::::::: ::::::::::::::::::::::::::::::::::::::::::::::::
P         AACGCAGGCGCUGCUCGGCGGCAUCUGGAGGCCAAGGGGUUCCCACUCGACGAGUUCCUA
                2050      2060      2070      2080      2090      2100

                2110      2120      2130      2140      2150      2160
WT        GCCGAGUGGUCUGAGCUGUCAGAGUUCGGUGAGGCCUUCGAAGGCUUCAAUAUCAAGCUG
          ::::::::::: ::::::::::::::::::::::::::::::::::::::::::::::::
P         GCCGAGUGGUCCGAGCUGUCAGAGUUCGGUGAGGCCUUCGAAGGCUUCAAUAUCAAGCUG
                2110      2120      2130      2140      2150      2160

                2170      2180      2190      2200      2210      2220
WT        ACCGUAACAUCUGAGAGCCUAGCCGAACUGAACAAGCCAGUACCCCCCAAGCCCCCAAAU
          ::::::::::::::::::::::::::::::::::::::::::::::::::::::::::::
P         ACCGUAACAUCUGAGAGCCUAGCCGAACUGAACAAGCCAGUACCCCCCAAGCCCCCAAAU
                2170      2180      2190      2200      2210      2220

                2230      2240      2250      2260      2270      2280
```

```
WT     GUCAACAGACCAGUCAACACUGGGGGACUCAAGGCAGUCAGCAACGCCCUCAAGACCGGU
       :::::::::::::::::::::::::::::::::::::::::::::::::: :::::::::
P      GUCAACAGACCAGUCAACACUGGGGGACUCAAGGCAGUCAGCAACGCCCUUAAGACCGGU
             2230      2240      2250      2260      2270      2280

             2290      2300      2310      2320      2330      2340
WT     CGGUACAGGAACGAAGCCGGACUGAGUGGUCUCGUCCUUCUAGCCACAGCAAGAAGCCGU
       ::::::::::::::::::::::::::::::::::::::::::::::::::::: ::::::
P      CGGUACAGGAACGAAGCCGGACUGAGUGGUCUCGUCCUUCUAGCCACAGCAAGGAGCCGU
             2290      2300      2310      2320      2330      2340

             2350      2360      2370      2380      2390      2400
WT     CUGCAAGAUGCAGUUAAGGCCAAGGCAGAAGCCGAGAAACUCCACAAGUCCAAGCCAGAC
       :::::::: :::::::::::::::::::::::::::::::::::::::::::::::::::
P      CUGCAAGACGCAGUUAAGGCCAAGGCAGAAGCCGAGAAACUCCACAAGUCCAAGCCAGAC
             2350      2360      2370      2380      2390      2400

             2410      2420      2430      2440      2450      2460
WT     GACCCCGAUGCAGACUGGUUCGAAAGAUCAGAAACUCUGUCAGACCUUCUGGAGAAAGCC
       :::::::::::::::::::: :::::::::::::::::::::::::::::::::::::::
P      GACCCCGAUGCAGACUGGUUUGAAAGAUCAGAAACUCUGUCAGACCUUCUGGAGAAAGCC
             2410      2420      2430      2440      2450      2460

             2470      2480      2490      2500      2510      2520
WT     GACAUCGCCAGCAAGGUCGCCCACUCAGCACUCGUGGAAACAAGCGACGCCCUUGAAGCA
       ::::::::::: ::::::::::::::::::::::::::::::::::::::: :::::::::
P      GACAUCGCCAGUAAGGUCGCCCACUCAGCACUCGUGGAAACAAGCGACGCUCUUGAAGCA
             2470      2480      2490      2500      2510      2520

             2530      2540      2550      2560      2570      2580
WT     GUUCAGUCGACUUCCGUGUACACCCCCAAGUACCCAGAAGUCAAGAACCCACAGACCGCC
       ::::::::::::::::::::::::::::::::::::::::::::::::::::::::::::
P      GUUCAGUCGACUUCCGUGUACACCCCCAAGUACCCAGAAGUCAAGAACCCACAGACCGCC
             2530      2540      2550      2560      2570      2580

             2590      2600      2610      2620      2630      2640
WT     UCCAACCCCGUUGUUGGGCUCCACCUGCCCGCCAAGAGAGCCACCGGUGUCCAGGCCGCU
       ::::::::::::::::::::::::::::::::::::::::::::::::::::::::::::
P      UCCAACCCCGUUGUUGGGCUCCACCUGCCCGCCAAGAGAGCCACCGGUGUCCAGGCCGCU
             2590      2600      2610      2620      2630      2640

             2650      2660      2670      2680      2690      2700
WT     CUUCUCGGAGCAGGAACGAGCAGACCAAUGGGGAUGGAGGCCCCAACACGGUCCAAGAAC
       ::::::::::::::::::::: ::::::::::::::::::::::::::::::::::::::
P      CUUCUCGGAGCAGGAACGAGCCGACCAAUGGGGAUGGAGGCCCCAACACGGUCCAAGAAC
             2650      2660      2670      2680      2690      2700

             2710      2720      2730      2740      2750      2760
WT     GCCGUGAAAAUGGCCAAACGGCGGCAACGCCAAAAGGAGAGCCGCUAACAGCCAUGAUGG
       ::::::::::::::::::::::::::::::::::: ::::::::: :: :::::::::::
P      GCCGUGAAAAUGGCCAAACGGCGGCAACGCCAAAAAGAGAGCCGCCAAUAGCCAUGAUGG
             2710      2720      2730      2740      2750      2760

             2770      2780      2790      2800      2810      2820
WT     GAACCACUCAAGAAGAGGACACUAAUCCCAGACCCCGUAUCCCCGGCCUUCGCCUGCGGG
       ::::::::::::::::::::::::::::::::::::::::::::::::::::::::::::
P      GAACCACUCAAGAAGAGGACACUAAUCCCAGACCCCGUAUCCCCGGCCUUCGCCUGCGGG
             2770      2780      2790      2800      2810      2820


WT     GGCCCCC
       :::::::
P      GGCCCCC

98.0% identity in 2,827 nt overlap
```

#### 3. Protein Comparisons

Comparisons were carried out with the SIM Alignment Tool for protein sequences (https://web.expasy.org/sim/)using default parameters.

#### 3.1. Non-structural Protein; VP5

```
WT      MVSRDQTNDRSDDKPARSNPTDCSVHTEPSDANNRTGVHSGRHPGEAHSQVRDLDLQFDC        60
P       MVSRDQTNDRSDDKPDRSNPTDCSVHTEPSDANNRTGVHSGRHPGEAHSQVRDLDLQFDC        60
        *************** ********************************************

WT      GGHRVRANCLFPWIPWLNCGCSLHTAGQWELQVRSDAPDCPEPTGQLQLLQASESESHSE        120
P       GGHRVRANCLFPWIPWLNCGCSLHTAEQWELQVRSDAPDCPEPTGQLQLLQASESESHSE        120
        ************************** *********************************

WT      VKHTSWWRLCTKRHHKRRDLPRKPE     145
P       VKHTPWWRLCTKRHHKRRDLPRKPE     145
        **** ********************
```

97.9% identity in 145 residues ovrelap

#### 3.2. Polyprotein

```
WT      MTNLQDQTQQIVPFIRSLLMPTTGPASIPDDTLEKHTLRSETSTYNLTVGDTGSGLIVFF        60
P       MTNLTDQTQQIVPFIRSLLMPTTGPASIPDDTLEKHTLRSETSTYNLTVGDTGSGLIVFF        60
        **** *******************************************************

WT      PGFPGSIVGAHYTLQGNGNYKFDQMLLTAQNLPASYNYCRLVSRSLTVRSSTLPGGVYAL        120
P       PGFPGSIVGAHYTLQSNGNYKFDQMLLTAQNLPASYNYCRLVSRSLTVRSSTLPGGVYAL        120
        ***************.********************************************

WT      NGTINAVTFQGSLSELTDVSYNGLMSATANINDKIGNVLVGEGVTVLSLPTSYDLGYVRL        180
P       NGTINAVTFQGSLSELTDVSYNGLMSATANINDKIGNVLVGEGVTVLSLPTSYDLGYVRL        180
        ************************************************************

WT      GDPIPAIGLDPKMVATCDSSDRPRVYTITAADDYQFSSQYQPGGVTITLFSANIDAITSL        240
P       GDPIPAIGLDPKMVATCDSSDRPRVYTITAADDYQFSSQYQSGGVTITLFSANIDAITSL        240
        ***************************************** ******************

WT      SVGGELVFRTSVHGLVLGATIYLIGFDGTTVITRAVAANNGLTTGTDNLMPFNLVIPTNE        300
P       SIGGELVFHTSVHGLALDATIYLIGFDGTTVITRAVASDNGLTTGIDNLMPFNLVIPTNE        300
        *:******:******.*.*******************::****** **************

WT      ITQPITSIKLEIVTSKSGGQAGDQMSWSARGSLAVTIHGGNYPGALRPVTLVAYERVATG        360
P       ITQPITSIKLEIVTSKSGGQAGDQMSWSASGSLAVTIHGGNYPGALRPVTLVAYERVATG        360
        ***************************** ******************************

WT      SVVTVAGVSNFELIPNPELAKNLVTEYGRFDPGAMNYTKLILSERDRLGIKTVWPTREYT        420
P       SVVTVAGVSNFELIPNPELAKNLVTEYGRFDPGAMNYTKLILSERDRLGIKTVWPTREYT        420
        ************************************************************

WT      DFREYFMEVADLNSPLKIAGAFGFKDIIRAIRRIAVPVVSTLFPPAAPLAHAIGEGVDYL        480
P       DFREYFMEVADLNSPLKIAGAFGFKDIIRAIRRIAVPVVSTLFPPAAPLALAIGEGVDYL        480
        ************************************************** *********

WT      LGDEDQAASGTARAASGKARAASGRIRQLTLAADKGYEVVANLFQVPQNPVVDGILASPG        540
P       LGDEAQAASGTARAASGKARAASGRIRQLTLAADKGYEVVANLFQVPQNPVVDGILASPG        540
        **** *******************************************************

WT      VLRGAHNLDCVLREGATLFPVVITTVEDAMTPKALNSKMFAVIEGVREDLQPPSQRGSFI        600
P       VLRGAHNLDCVLREGATLFPVVITTVEDAMTPKALNSKMFAVIEGVREDLQPPSQRGSFI        600
        ************************************************************

WT      RTLSGHRVYGYAPDGVLPLETGRDYTVVPIDDVWDDSIMLSKDPIPPIVGNSGNLAIAYM        660
P       RTLSGHRVYGYAPDGVLPLETGRDYTVVPIDDVWDDSIMLSKDPIPPIVGNSGNLAIAYM        660
        ************************************************************

WT      DVFRPKVPIHVAMTGALNACGEIEKVSFRSTKLATAHRLGLRLAGPGAFDVNTGPNWATF        720
P       DVFRPKVPIHVAMTGALNACGEIEKVSFRSTKLATAHRLGLKLAGPGAFDVNTGPNWATF        720
        *****************************************:******************

WT      IKRFPHNPRDWDRLPYLNLPYLPPNAGRQYHLAMAASEFKETPELESAVRAMEAAANVDP        780
P       IKRFPHNPRDWDRLPYLNLPYLPPNAGRQYHLAMAASEFKETPELESAVRAMEAAANVDP        780
        ************************************************************

WT      LFQSALSVFMWLEENGIVTDMANFALSDPNAHRMRNFLANAPQAGSKSQRAKYGTAGYGV        840
P       LFQSAISVFMWLEENGIVTDMANFALSDPNAHRMRNFLANAPQAGSKSQRAKYGTAGYGV        840
```

```
         *****:******************************************************

WT       EARGPTPEEAQREKDTRISKKMETMGIYFATPEWVALNGHRGPSPGQLKYWQNTREIPDP         900
P        EARGPTPEEAQREKDTRISKKMETMGIYFATPEWVALNGHRGPSPGQLKYWQNTREIPDP         900
         ************************************************************

WT       NEDYLDYVHAEKSRLASEEQILRAATSIYGAPGQAEPPQAFIDEVAKVYEINHGRGPNQE         960
P        NEDYLDYVHAEKSRLASEEQIQRAATSIYGAPGQAEPPQAFIDEVAKVYEINHGRGPNQE         960
         ********************* **************************************

WT       QMKDLLLTAMEMKHRNPRRALPKPKPKPNAPTQRPPGRLGRWIRTVSDEDLE       1012
P        QMKDLLLTAMEMKHRNPRRAPPKPKPKPNAPTPRPPGRLGRWIRTVSDEDLE       1012
         ******************** *********** *******************
```

98.2% identity in 1,012 residues overlap

### 3.3. RNA Dependent RNA Polymerase; VP1

```
WT       MSDVFNSPQARSTISAAFGIKPTAGQDVEELLIPKVWVPPEDPLASPSRLAKFLRENGYK         60
P        MSDIFNSPQARSKISAAFGIKPTAGQDVEELLIPKVWVPPEDPLASPSRLAKFLRENGYK         60
         ***:********.***********************************************

WT       VLQPRSLPENEEYETDQILPDLAWMRQIEGAVLKPTLSLPIGDQEYFPKYYPTHRPSKEK         120
P        VLQPRSLPENEEYETDQILPDLAWMRQIEGAVLKPTLSLPIGDQEYFPKYYPTHRPSKEK         120
         ************************************************************

WT       PNAYPPDIALLKQMIYLFLQVPEANEGLKDEVTLLTQNIRDKAYGSGTYMGQATRLVAMK         180
P        TNAYPPDIALLKQMIYLFLQVPEANEGLKDEVTLLTQNIRDKAYGSGTYMGQATRLVAMK         180
          ***********************************************************

WT       EVATGRNPNKDPLKLGYTFESIAQLLDITLPVGPPGEDDKPWVPLTRVPSRMLVLTGDVD         240
P        EVATGRNPNKDPLKLGYTFESIAQLLDITLPVGPPGEDDKPWVPLTRVPSRMLVLTGDVD         240
         ************************************************************

WT       GDFEVEDYLPKINLKSSSGLPYVGRTKGETIGEMIAISNQFLRELSTLLKQGAGTKGSNK         300
P        GDFEVEDYLPKINLKSSSGLPYVGRTKGETIGEMIAISNQFLRELSTLLKQGAGTKGSNK         300
         ************************************************************

WT       KKLLSMLSDYWYLSCGLLFPKAERYDKSTWLTKTRNIWSAPSPTHLMISMITWPVMSNSP         360
P        KKLLSMLSDYWYLSCGLLFPKAERYDKSTWLTKTRNIWSAPSPTHLMISMITWPVMSNSP         360
         ************************************************************

WT       NNVLNIEGCPSLYKFNPFRGGLNRIVEWILAPEEPKALVYADNIYIVHSNTWYSIDLEKG         420
P        NNVLNIEGCPSLYKFNPFRGGLNRIVEWILAPEEPKALVYADNIYIVHSNTWYSIDLEKG         420
         ************************************************************

WT       EANCTRQHMQAAMYYILTRGWSDNGDPMFNQTWATFAMNIAPALVVDSSCLIMNLQIKTY         480
P        EANCTRQHMQAAMYYILTRGWSDNGDPMFNQTWATFAMNIAPALVVDSSCLIMNLQIKTY         480
         ************************************************************

WT       GQGSGNAATFINNHLLSTLVLDQWNLMRQPRPDSEEFKSIEDKLGINFKIERSIDDIRGK         540
P        GQGSGNAATFINNHLLSTLVLDQWNLMRQPRPDSEEFKSIEDKLGINFKIERSIDDIRGK         540
         ************************************************************

WT       LRQLVLLAQPGYLSGGVEPEQSSPTVELDLLGWSATYSKDLGIYVPVLDKERLFCSAAYP         600
P        LRQLVPLAQPGYLSGGVEPEQSSPTVELDLLGWSATYSKDLGIYVPVLDKERLFCSAAYP         600
         ***** ******************************************************

WT       KGVENKSLKSKVGIEQAYKVVRYEALRLVGGWNYPLLNKACKNNAGAARRHLEAKGFPLD         660
P        KGVENKSLKSKVGIEQAYKVVRYEALRLVGGWNYPLLNKACKNNAGAARRHLEAKGFPLD         660
         ************************************************************

WT       EFLAEWSELSEFGEAFEGFNIKLTVTSESLAELNKPVPPKPPNVNRPVNTGGLKAVSNAL         720
P        EFLAEWSELSEFGEAFEGFNIKLTVTSESLAELNKPVPPKPPNVNRPVNTGGLKAVSNAL         720
         ************************************************************

WT       KTGRYRNEAGLSGLVLLATARSRLQDAVKAKAEAEKLHKSKPDDPDADWFERSETLSDLL         780
P        KTGRYRNEAGLSGLVLLATARSRLQDAVKAKAEAEKLHKSKPDDPDADWFERSETLSDLL         780
         ************************************************************

WT       EKADIASKVAHSALVETSDALEAVQSTSVYTPKYPEVKNPQTASNPVVGLHLPAKRATGV         840
P        EKADIASKVAHSALVETSDALEAVQSTSVYTPKYPEVKNPQTASNPVVGLHLPAKRATGV         840
         ************************************************************
```

```
WT      QAALLGAGTSRPMGMEAPTRSKNAVKMAKRRQRQKESR- 878
P       QAALLGAGTSRPMGMEAPTRSKNAVKMAKRRQRQKESRQ 879
        **************************************
```

99.5% identity in 878 residues overlap
